# The use of cultured symbionts enhances the survival of reef-deployed coral recruits

**DOI:** 10.64898/2026.11.14.766699

**Authors:** E Pfeffer, MR Nitschke, S Ferguson, Y Sato, MA Abdul Wahab

## Abstract

**Introduction:** Symbiodiniaceae dinoflagellates are essential for coral juvenile development and survival, yet early-life symbiont provisioning remains underused in coral restoration.

**Objectives:** We tested whether inoculating sexually propagated *Acropora spathulata* recruits with cultured symbionts before reef deployment improves survival and shapes symbiont community composition.

**Methods:** Recruits inoculated with cultured *Cladocopium* sp. (C1) or *Durusdinium trenchii* (D1), and uninoculated recruits (Control) were deployed on the central Great Barrier Reef.

**Results:** Both inoculation treatments approximately doubled survival relative to Control, although the benefit varied with local benthic habitat. C1 and D1 recruits were dominated by symbiont sequences matching their respective inoculated cultures, while Control recruits predominantly harboured a *Durusdinium*-dominant community closely resembling the cultured D1. Cultured C1 was also dominant among most C1-inoculated recruits after nine months, although some diverged from the inoculated community towards a mixed assemblage with *Durusdinium*.

**Conclusions:** Similar survival benefits in the two inoculation treatments suggest that the timing and success of symbiont establishment were more critical for early survival than symbiont genus identity, under the conditions and period of this study.

**Implications for Practice:**

- Inoculating coral recruits with symbionts before deployment improved early survival and could be incorporated into existing sexual propagation and coral-seeding workflows.
- Comparable survival of deployed recruits inoculated with *Cladocopium* or *Durusdinium* suggests that practitioners can prioritise symbionts that establish early and successfully over a particular genus.
- Benefits varied with local benthic context, so deployment sites and natural symbiont communities should be considered alongside symbiont provisioning.
- Future work should test whether the survival benefit holds across species, reefs, and thermal conditions, including under heat stress.

## INTRODUCTION

Marine heatwaves are increasing in frequency and severity because of global climate change, driving widespread coral bleaching and declines in coral cover worldwide (Hughes et al. 2018). While reducing greenhouse gas emissions remains essential, active restoration has grown substantially as a means of enhancing reef recovery and resilience. Larval culture and deployment of coral recruits has emerged as a scalable restoration approach, with the potential to maintain genetic diversity generated through sexual reproduction (Bayraktarov et al. 2016; Boström-Einarsson et al. 2020; Dallmeyer-Drennen et al. 2026). This typically involves settlement of larvae onto engineered substrates or seeding devices followed by deployment to reefs. Under typical conditions, more than 90% of newly settled corals die within the first six months on the reef (Vermeij & Sandin 2008; Penin et al. 2010), with mortality driven by competition with benthic organisms, sedimentation, predation, grazing pressure, and energetic limitation (Kuffner et al. 2006; Page et al. 2024; Humanes et al. 2025). Despite substantial recent progress in gamete harvesting (Doropoulos et al. 2019b) and larval culturing (Doropoulos et al. 2019a; Severati et al. 2024; Ramsby et al. 2026a), seeding device design, deployment methodology, and operational efficiency (Chamberland et al. 2017; Mendoza Quiroz et al. 2025; Ramsby et al. 2026b), post-deployment survival remains a key constraint on scaling larval-based restoration (Page et al. 2024; Jurriaans et al. 2025; Whitman et al. 2025).

As reef-building corals function as holobionts, post-settlement survival can increase with the establishment of symbiosis with dinoflagellates of the family Symbiodiniaceae, which in adult colonies provide the host’s daily metabolic energy through photosynthetically derived carbon (Muscatine & Porter 1977; Yellowlees et al. 2008). Most coral larvae are initially aposymbiotic and rely on maternally provisioned lipid reserves but progressively transition towards symbiont-derived nutrition as settlement and calcification proceed (Huffmyer et al. 2025). Approximately 80–85% of coral species acquire symbionts from the environment (Baird et al. 2009; Quigley et al. 2017); acquisition depends on environmental availability, host selectivity, and symbiont infectivity (Abrego et al. 2009a, 2009b; 2012).

Symbiodiniaceae are highly diverse (LaJeunesse et al. 2018), and their identity strongly influences coral physiology and performance, including growth, survival, thermal tolerance, and calcification at both adult and early life stages (Baker 2003; Little et al. 2004; Quigley et al. 2018). Notably, *Durusdinium* symbionts are commonly associated with increased thermal tolerance but may impose physiological trade-offs, whereas *Cladocopium* symbionts are often linked to higher growth under favourable conditions (Berkelmans & van Oppen 2006; Jones & Berkelmans 2010; Cunning et al. 2015). Selective symbiont provisioning may therefore provide a means to improve restoration outcomes. However, existing field data address either recruit performance without characterising the resulting symbiont communities (Waters et al. 2025; Whitman et al. 2025; Jurriaans et al. 2026) or symbiont uptake following inoculation with strains freshly isolated from conspecific adult colonies, assessed under laboratory or short-term conditions rather than following field deployment (Little et al. 2004; Abrego et al. 2012). To our knowledge, no prior field study has paired survival outcomes with symbiont community characterisation in device-deployed, sexually propagated coral recruits, leaving it unknown whether the identity of inoculated symbionts contributes to early post-deployment performance. It was also untested whether targeted inoculation with strains from established culture collections, prior to deployment, could enhance the survival outcomes of recruits under specific reef-relevant conditions.

This study tested whether symbiont inoculation can enhance survival and shape symbiont composition in sexually propagated *Acropora spathulata* recruits deployed on a mid-shelf reef of the Great Barrier Reef (GBR). Specifically, we assessed whether inoculation with cultured *Cladocopium* sp. (hereafter ‘C1’) or *Durusdinium trenchii* (hereafter ‘D1’) (1) improves early post-deployment survival relative to non-inoculated controls, and (2) the inoculum remains detectable in recruits sampled at three and nine months despite exposure to environmental symbiont pools and natural reef conditions.

## METHODS

### Coral production

Twelve gravid colonies of *Acropora spathulata* (∼20 cm diameter) were collected using a hammer and chisel on SCUBA at depths of 2−5 m from Davies Reef, central GBR, Australia (18° 49’ 12.2’’ S, 147° 38’ 39.4’’ E), on 27 November 2023 (full moon). Gravid colonies were determined by visual inspection of pigmented eggs within the coral tissue *in situ*. Fragments were transported in 70 L bins (1−4 fragments per bin) receiving flow-through seawater onboard a research vessel over a period of 4−6h to the National Sea Simulator (Australian Institute of Marine Science, AIMS, Townsville). Corals were held in outdoor semi-recirculating aquaria that received new input of 1µm filtered seawater (FSW) at a rate of ∼3 turnovers per day and that profiled the ambient temperature experienced at Davies Reef (∼27.2°C).

On 30 November, upon detection of setting, colonies were isolated in individual 70 L bins for spawning. Upon release (∼21:25−22:50 h), buoyant egg-sperm bundles from each colony were collected by skimming the water surface using a plastic cup. Bundles were then added to a single Auto Fertiliser module (85 L; see Severati et al. 2024 for system design and procedure), which achieved 80 ± 0.08 (SE) % fertilisation success after 1.5 h. Fertilised eggs were stocked (0.5−0.9 larva mL^-1^) across 5 × 500L culture tanks receiving flow-through filtered sea water (FSW) at an exchange rate of 20% hr^-1^ following Ramsby et al. 2026a. Culture tanks received low irradiance from fluorescent lighting in the facility (∼5 µmol m^-2^ s^-1^). Larvae were reared over a period of 7 days. Once competency for settlement of >75% was reached, larvae were mass settled onto concrete tiles (280 × 280 mm) over a period of 24−48 h (Ramsby et al. 2026a). Concrete tiles were conditioned in outdoor tanks receiving seawater over a period of 9 weeks to develop microbial biofilm and crustose coralline algae to induce settlement. Prior to settlement, tiles were frozen for ∼48h at −20°C to kill any biota and flushed for 4h in clean FSW prior to use.

### Symbiodiniaceae culturing

Symbiodiniaceae cultures were maintained at the AIMS Symbiont Culture Facility (SCF). The C1 culture (SCF049.1), originally isolated from *Pocillopora damicornis*, and the D1 culture (SCF086.1), isolated from *Porites lobata*, were grown in sterile IMK for marine microalgae (Nihon Pharmaceutical Co., Ltd.). Cultures were maintained at 27°C under approximately 60 µmol photons m^⁻2^ s⁻¹ on a 14:10 hr light:dark photoperiod in temperature-controlled incubation chambers. Both cultures originated from Davies Reef and had been maintained in the SCF collection for 15 years.

To produce sufficient biomass for inoculation, cultures were progressively upscaled over three months prior to coral spawning through sequential subculturing in progressively larger sterile, single use vessels (T-25, T-75, T-175 culture flasks, 2 L bottles, 10 L carboys). Culture identity and purity in the SCF is routinely monitored using ITS2 metabarcoding, with the entire SCF culture collection genotyped approximately one to two times per year.

### Symbiont inoculation

Tiles (n = 2 per tank) with settled corals (termed ‘spat’ hereafter) were placed into 3 × 50 L flow-through tanks and inoculated with one of the following treatments: C1, D1, or a no-inoculation control. Flow-through was turned off to tanks during inoculation, each symbiont culture was added to achieve a final cell density of 1 × 10^4^ cells mL^-1^ and incubated for 12 hours overnight with the temperature maintained in a temperature-controlled room (∼27°C). Post-inoculation, flow-through FSW to the tanks was switched back on at 0.8L min^-1^(100% exchange hr^-1^) flow rate. Spat were held in the same tanks for 2 weeks under a 12:12 hr light cycle (40 µmol photons m^⁻2^ s⁻¹) and fed daily with rotifers (5 rotifer mL ^-1^), while awaiting device assembly.

### Device assembly and field deployment

After 2 weeks, settlement tiles were cut into individual tabs (14 × 14 mm), and tabs containing 3–5 coral spat selected and inserted into ceramic coral seeding devices. Each device held 3 tabs (Ramsby et al. 2026b); therefore, each device had between 9−15 individual coral spat. Devices were assembled by treatment groups, totaling 90 devices each for D1 and Control treatments, and 180 for C1 treatment. The assembly of each treatment was handled separately (isolated seawater sources) to prevent symbiont cross-contamination. However, all devices were stacked together vertically on stainless steel rods and placed into 2000 L semi-recirculating holding tanks for one week prior to deployment, without treatment distinction. At this point, treatments were exposed to each other. This was necessary to prepare devices for transport to the field.

A total of 360 seeding devices were deployed by SCUBA on the north-west flank of Davies Reef (18.80543°S, 147.64923°E) between the 28 to the 30 December 2023, at 7−9 m depth. Devices were distributed across fifteen 1m^2^ plots spaced approximately 4 m apart along a depth contour and marked with a central metal stake. Devices were designed with a ceramic linking pin to enable devices to be connected in chains (**Figure 1**), and central devices were fitted with an RFID tag (Biomark; Merck & Co., Inc., Rahway, NJ, USA) to enable identification of treatment chains if movement occurred within plots. Each plot contained four treatment chains comprising two chains of six C1-inoculated devices (12 devices), one chain of six D1-inoculated devices, and one chain of six non-inoculated Control devices (**Figure 1A**). Additional devices associated with other independent and unrelated experiments were deployed within plots but were not included in the present study (**Figure 1A**).

**Figure 1.**
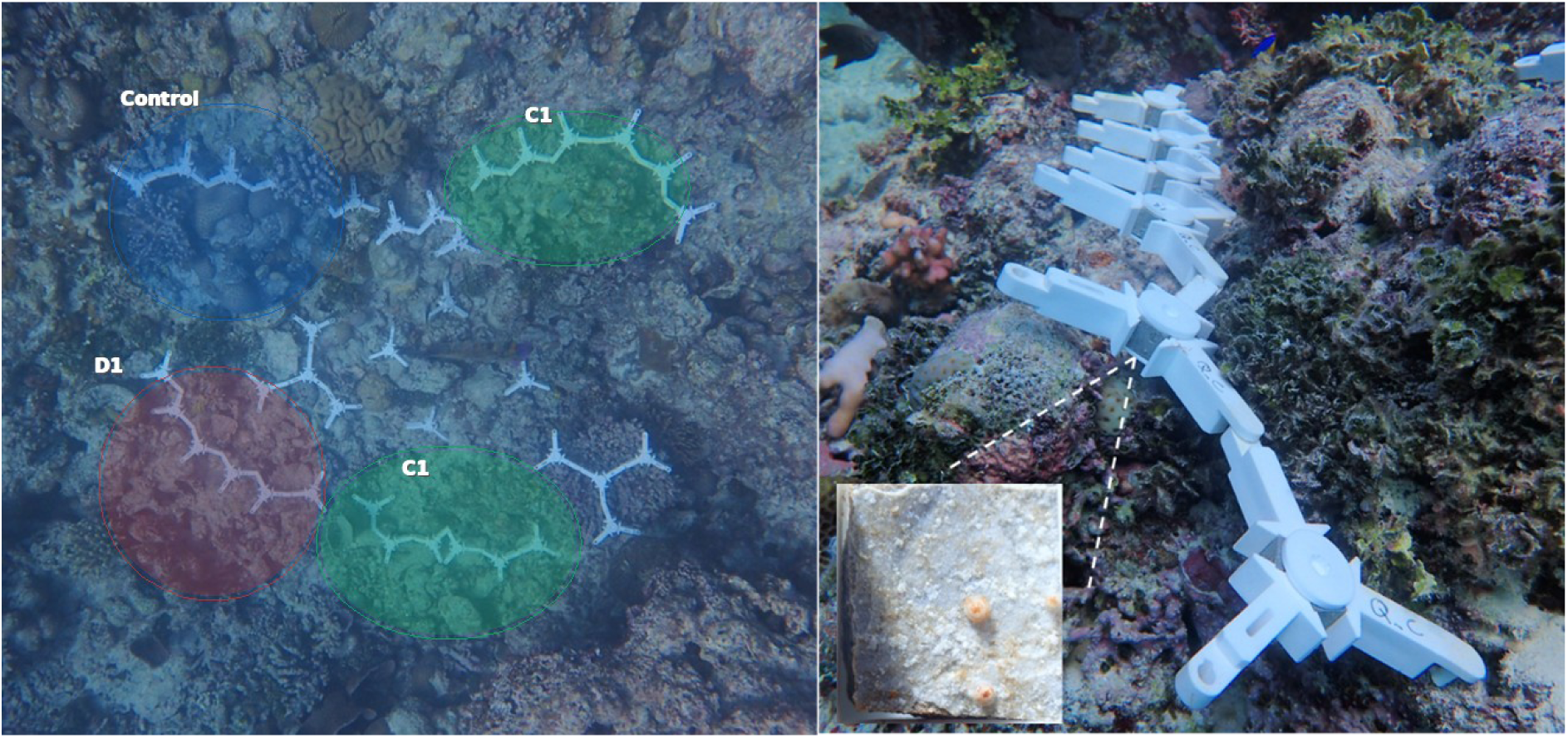
Experimental design and coral seeding devices at Davies Reef, central Great Barrier Reef. (A) Layout at each reef plot: seeding devices were linked in chains of six, with one chain of six Control devices (uninoculated recruits), one chain of six D1 devices (recruits inoculated with *Durusdinium trenchii*, strain SCF086) and two chains of six C1 devices (recruits inoculated with *Cladocopium* sp., strain SCF049; 12 devices). Chains were placed around a central steel reinforcing bar (rebar), alongside other independent experiments in the middle of the plot. (B) A representative chain of devices placed over the benthos. The inset shows newly settled coral spat on a tab, the unit on which survival was scored.

To characterise the site’s environmental conditions, a temperature logger (HOBO Water Temp Pro v2, Onset) was installed to measure temperature at 10-minute intervals. Additionally, the flow of seawater was captured using a Marotte HS drag-tilt current meter (James Cook University) measuring velocity every minute. Instruments were retrieved and redeployed at three-month intervals for data download, maintenance, and biofouling removal. Benthic composition at each plot was captured using an Olympus Tough TG-6 camera (Olympus Imaging America Inc), within a 50 × 50 cm quadrat placed in 4 directions (NW, NE, SE, SW) relative to the central metal picket. Images were manually annotated using ReefCloud (AIMS, 2024), characterising 20 random points per image to estimate percent cover of crustose coralline algae (CCA), macroalgae, sand, sediment, soft coral, hard coral, cyanobacteria, epilithic algal matrix (EAM), sponges, and other organisms.

### Survival assessment and sampling

Deployed spat (hereafter ‘recruits’) were assessed for survivorship and symbiont assemblage at three months post-deployment across all treatments. Because sampling was destructive, all D1 and Control devices (90 devices each) were retrieved at three months to ensure sufficient sample size. C1 devices (n = 180) were expected to retain sufficient surviving recruits for a further sampling point, and 90 devices remained deployed and were sampled after nine months. Survivorship was recorded across 15 plots for Control, and 14 plots for C1 and D1 treatments, from 210 settlement devices (83 Control, 44 C1-inoculated, 83 D1-inoculated) and 630 tabs (249 Control, 132 C1-inoculated, 249 D1-inoculated). Survivorship was quantified as the presence or absence of at least one living recruit on a tab, defined as yield. Tab-level yield was analysed using generalised linear mixed-effects models (GLMMs) with binomial error distribution and logit link. The primary model included treatment (Control, C1, D1) as the sole fixed effect of recruit survival. Plot and device identity were included as random intercepts to account for spatial clustering and non-independence among tabs originating from the same device.

To evaluate whether benthic taxa categories within plots modulated treatment effects on survival, we fitted a series of separate GLMMs, each testing the interaction between treatment and a single habitat predictor. Benthic habitat cover was quantified at the plot level (photoquadrats averaged to a single percent-cover value per benthic category per plot), each plot-level habitat value was therefore shared across all devices and tabs nested within that plot. Habitat covariates were standardised (z-scores) prior to fitting, and both plot and device identity were included as random intercepts to account for non-independence among tabs sharing the same plot-level habitat value and device. Fixed effects were tested using Type II Wald χ^2^ tests, and predicted marginal means were estimated for each model.

### DNA extraction and ITS2 community processing

After three months for all treatments, and after nine months for C1, individual recruits were sampled using sterile scalpels and preserved in 1 mL absolute ethanol. Instruments were sterilised between samples using a 3-step protocol: absolute ethanol, followed by 30% Sodium hypochlorite, then filtered seawater. Where multiple recruits were present on a single tab, individuals were sampled separately. Samples were stored at 4°C, ethanol replaced after 24 h, and samples transported to AIMS for storage at −20°C while awaiting molecular processing. For DNA extraction and ITS2 metabarcoding, we used recruits from C1 and Control treatments across 13 and 12 plots respectively, and D1 recruits from five plots, due to field sample availability at collection. Total DNA was extracted from recruits sampled using the Maxwell® RSC PureFood GMO and Authentication Kit (Promega) on the Maxwell® robot, following the manufacturer’s manual. Tissue fragments (<2 mm^2^) were lysed overnight in CTAB buffer at 56°C, and 300 µL lysate was combined with cell lysis buffer prior to automated extraction. DNA was eluted in a final volume of 100 µL of TE buffer. Extraction quality and concentration were assessed using Nanodrop spectrophotometry, Qubit dsDNA high-sensitivity assays, and agarose gel electrophoresis. Symbiodiniaceae communities were characterised using ITS2 amplicon sequencing. Libraries were prepared using a two-step PCR protocol consisting of an initial amplification with the SYM_VAR_5.8S2/SYM_VAR_REV primer pair (Hume et al. 2018), targeting the ITS2 region followed by a second indexing PCR to add Illumina sequencing adapters and dual indices. Each library included negative controls (PCR blanks and extraction controls) and included four positive culture controls corresponding to the two inoculated symbiont strains. Finally, indexed amplicon libraries were pooled together and sequenced on an Illumina NextSeq platform (Walter and Eliza Hall Institute). Raw sequencing reads received were demultiplexed using Cutadapt (v5.1). Demultiplexed FASTA files were submitted to SymPortal for ITS2 intragenomic variant analysis (Hume et al. 2019). Samples exhibiting read depth lower than 1500 reads were excluded and sequences detected in library negative controls were removed from corresponding samples.

### Effects of inoculation on symbiont genus composition

Symbiodiniaceae community composition and inoculum retention across treatments were first analysed at the genus level using the sequence reads assigned to specific genera by SymPortal. For each sample, ITS2 sequence reads were summed by genus and converted to read-based relative abundances (not corrected for ITS2 copy-number variation among genera (Davies et al. 2023)). Samples in which neither *Durusdinium* nor *Cladocopium* were detected, were assigned a relative abundance of zero for that genus, so that comparisons between genera and treatments were made across a consistent set of samples. Genus-level analyses focused on *Cladocopium* and *Durusdinium*, the two genera used for inoculation, as together they accounted for the large majority of ITS2 reads across samples, with other genera occurring at consistently low relative abundance and were not modelled individually (**Table S4A; Figure S3**).

Two beta regression models with a logit link were fitted for *Cladocopium* and *Durusdinium* relative abundance within samples, with plot identity as a random intercept to account for spatial clustering among samples collected within reef plots. The first model tested the effect of inoculation treatment (Control, C1, D1) on *Cladocopium* and *Durusdinium* composition three months after deployment. The second compared the composition of these genera between recruits sampled at three and nine months post-deployment within C1-inoculated recruits only, since Control and D1 devices were fully retrieved at three months; timepoint was therefore fitted as the sole fixed effect in this model. Relative abundances of exactly 0 or 1 were adjusted by a small constant (0.001) prior to fitting, as required by the beta distribution’s open (0,1) interval. For both sets of models, estimates were back-transformed from the logit scale to the response scale to obtain predicted relative abundances and associated 95% confidence intervals. Pairwise comparisons for the first model were adjusted for multiple testing using the Tukey method and the single planned contrast between three and nine months within C1 required no such adjustment. Beta models showed departures from residual uniformity, reflecting the bimodal distribution of relative abundance among recruits, results are therefore interpreted alongside effect sizes and the fingerprint analyses.

### Culture-specific fingerprint construction and ITS2 sequence similarity analyses

Field validation of symbiont inoculum stability benefits from distinguishing cultured strains from environmentally acquired conspecific symbionts. SymPortal (Hume et al. 2019) resolves Symbiodiniaceae composition at the level of defining intragenomic sequence variants (DIVs), assigning ITS2 type profiles from sets of DIVs that recur identically across samples. This depends on separating intragenomic from intergenomic variation, which becomes harder in complex communities with few replicates or low read depth (Davies et al. 2023). Applied to our field dataset, this exact-match approach generated many distinct profiles per genus (33 for *Durusdinium*, 69 for *Cladocopium*), most sharing the same dominant DIVs as the cultured inoculum but differing in a variable tail of low-abundance accessory DIVs unique to individual samples. This precluded direct comparison of field samples to a single culture-derived profile. We therefore built culture fingerprint profiles directly from pure culture DIV sequences, which give cleaner baselines than host tissue containing multiple genera, species or strains (Davies et al. 2023) and allow consistent comparison of the culture-defining sequences across all field samples, independent of this background variation. Fingerprints were built for the C1 (SCF049) and D1 (SCF086) cultures sequenced alongside field samples in the same Illumina NextSeq library and subsequent SymPortal analysis. Sequences were ranked by median relative abundance across available culture replicates (two for SCF086; one only for SCF049, after a second replicate was removed during data cleaning; **Table S5A**).

Each fingerprint was defined as the set of top-ranked sequences whose cumulative median relative abundance reached 98% of the culture’s total signal, retaining 12 sequences for SCF049 and 8 for SCF086. Fingerprint profiles were then built from the median relative abundance of each fingerprint sequence across replicates, renormalised to sum to one.

Because ITS2 profiles are most informative when both the assemblage of sequences and their relative abundances are considered (Davies et al. 2023), retention of the inoculated symbionts was assessed with two complementary metrics: (1) fingerprint contribution, the proportion of a sample’s total ITS2 reads assigned to the same fingerprint sequences, and (2) fingerprint similarity, which quantifies how closely the abundance pattern of those contributing sequences matches the culture’s fingerprint profile. For each sample, fingerprint-sequences were renormalised to sum to one within fingerprint space, matching the fingerprint profile’s normalisation, to allow comparison to the fingerprint profile. Fingerprint similarity was calculated as 1 – Bray Curtis dissimilarity (BC), and a similarity threshold of 0.95 was chosen to indicate that the inoculum was retained within a sample, even where it represented only a subset of that sample’s total symbiont population. All statistical analyses were conducted in R version 4.6.0 (R Core Team 2026), using glmmTMB, car, DHARMa and emmeans (**Figures S2, S4, Tables S2C, S4H**). Analyses integrated packages from the tidyverse ecosystem and SymPortal outputs.

## RESULTS

### Symbiont inoculation increased recruit survival

Environmental conditions at Davies Reef were monitored from January to April 2024 (**Figure S1**). The monthly mean temperature decreased by approximately 0.8°C by April (January, 29.33 ± 0.43°C [mean ± SD]; April, 28.53 ± 0.68°C). Monthly current speed across the whole period did not exceed 0.04 m s⁻¹, apart from a short-lived peak in current speed in late January (daily mean ≈ 0.15 m s⁻¹; **Figure S1**) that coincided with a ∼0.8°C cooling, followed by the warmest period of the deployment in early February (daily mean ∼30°C).

After the first three months of the experiment, yield varied strongly across symbiont provisioning treatments. Observed tab – level yield after three months was 31.0% ± 5.4 (SE) in the Control group, 63.5% ± 4.4 in C1-inoculated, and 57.1% ± 7.1 in D1-inoculated recruits (**Figure 2**, **Table S1A**). Treatment had a significant effect on yield (Type II Wald χ^2^_2_ = 31.99, p < 0.001; 630 tabs on 210 devices in 15 plots; **Table S1B, S1E).** Model-predicted yield followed the observed pattern, whereby the C1- and D1-inoculated treatments were 2.6-fold (67.2%; 52.2–79.4) and 2.3-fold (60.0%; 47.4–71.5) higher than Control (**Table S1C**). Both inoculated treatments significantly outperformed Control (Tukey-adjusted p < 0.001), with no significant difference was detected between C1 and D1 (p = 0.630, **Table S1D**). Variation among devices within plots (variance = 1.26, SD = 1.12), exceeded variation among plots (variance = 0.42, SD = 0.65; **Table S1E**), indicating that fine-scale differences contributed more to variation in recruit survival than differences between plots.

**Figure 2.**
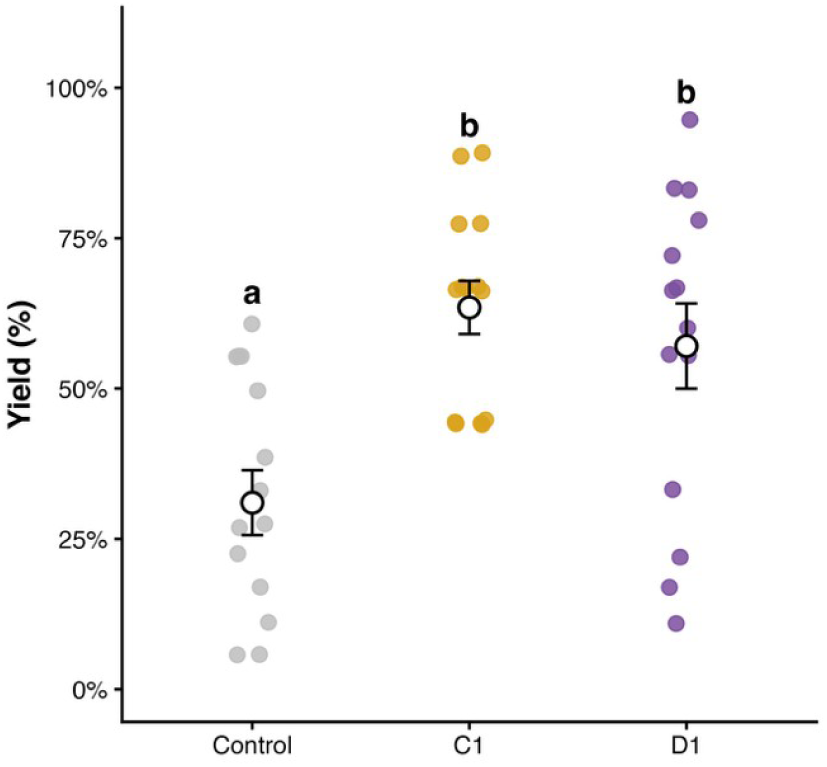
Observed yield (survival) of *Acropora spathulata* recruits three months after deployment, by treatment. Coloured dots show the mean yield for each reef plot (Control n = 15, C1 n = 14, D1 n = 14 plots). Open circles and error bars show the treatment mean ±1 SE across plots. Control, uninoculated recruits; C1, *Cladocopium* sp. SCF049; D1, *Durusdinium trenchii* SCF086. Treatments sharing a letter do not differ (Tukey-adjusted pairwise comparisons from a binomial generalised linear mixed model, p < 0.05).

Benthic cover was dominated by EAM cover (mean 72.8%, range 57.9–87.0%), followed by CCA (mean 10.4%, range 1.4–33.1%); all other categories averaged below 6% (**Table S2A**). Habitat cover alone showed no significant relationship with yield (all habitat main effects p > 0.05; **Table S2B**), but significant interactions between treatment and sediment, sand, CCA, soft coral, and cyanobacteria cover were detected (**Figure 3**). The effect of inoculation therefore varied with local benthic context, and the two symbionts responded in opposite directions: D1 recruits had higher predicted yield in plots with more CCA and less sediment, sand, and soft coral, C1 recruits showed the opposite pattern, and Control yield changed little across the sediment, sand, and soft coral gradients (**Figure 3; Table S2B**). Effect sizes (the change in predicted yield across a habitat’s observed range) were large relative to the narrow ranges over which sediment, sand, and soft coral varied (**Table S2A–B**), and C1’s response to CCA exceeded D1’s in magnitude. Cyanobacteria showed a similar contrast: predicted yield increased in Control and D1 recruits but declined in C1 recruits (**Figure 3**).

**Figure 3.**
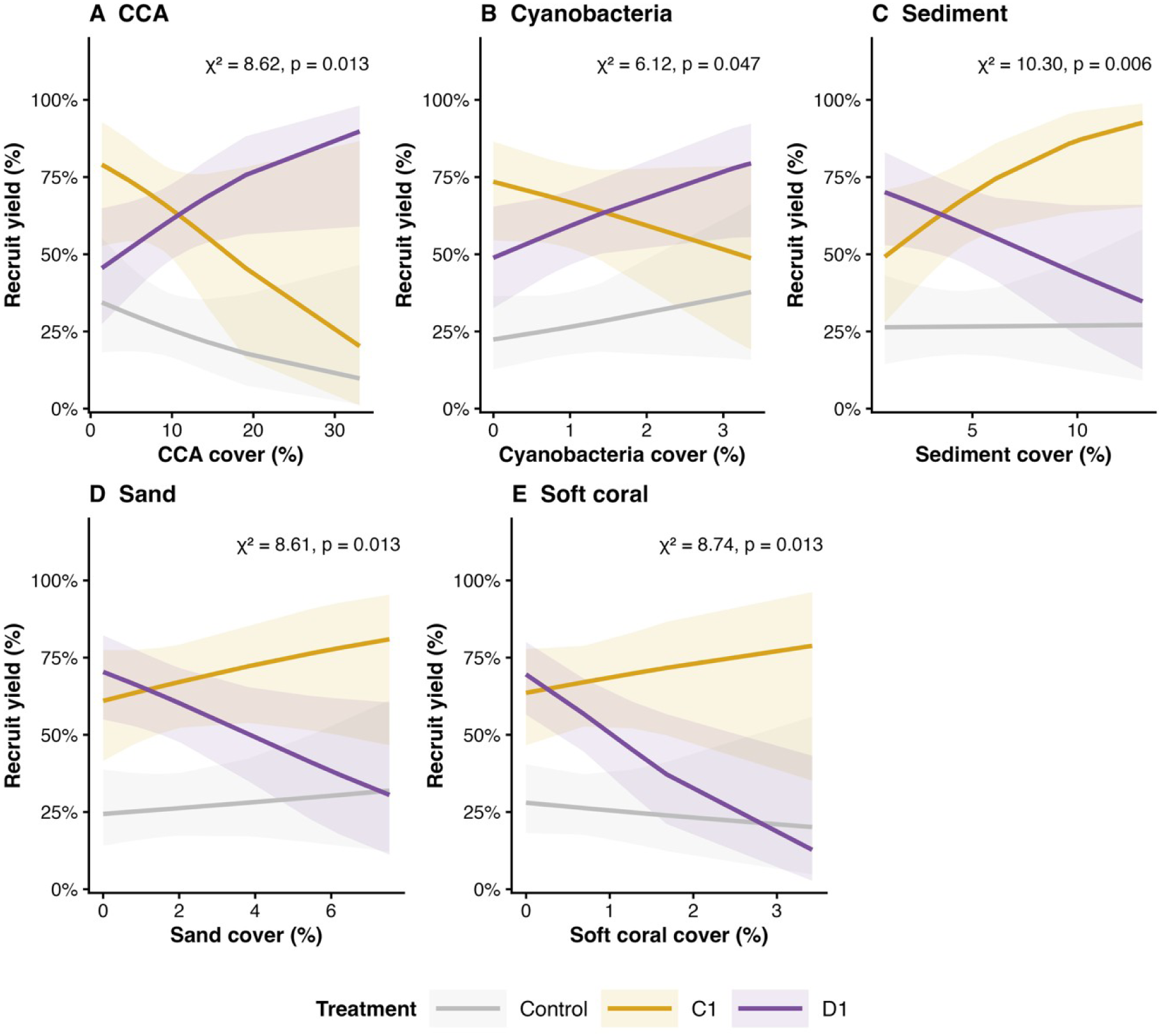
Model-predicted tab-level yield across benthic habitat gradients, by treatment. Each panel shows predicted yield as a function of plot-level cover of one benthic category (CCA = crustose coralline algae), fitted separately for Control, C1 and D1 recruits. Predictions come from a separate binomial generalised linear mixed model (GLMM) for each habitat, including treatment, habitat cover and their interaction, with plot and device as random intercepts. Shaded ribbons show 95% confidence intervals. Annotations in each panel give the Type II Wald χ^2^ and p-value for the treatment × habitat interaction.

### Symbiont inoculation shaped the symbiont composition of recruits

A total of 366 samples successfully passed the sequencing and cleaning process (3 months: Control, n = 71; C1-inoculated, n = 91; D1-inoculated, n = 87; 9 months: C1-inoculated, n = 117; **Table S3**). After three months on the reef, the relative abundance of *Cladocopium* and *Durusdinium* differed significantly among treatments (**Figure 4A–B**). C1-inoculated recruits were dominated by *Cladocopium*, whereas Control and D1-inoculated recruits were both dominated by *Durusdinium*. Treatment had a significant effect on *Cladocopium* relative abundance (Wald χ^2^₂ = 422.69, p < 0.001; **Table S4C**), with model-predicted proportions highest in C1-inoculated recruits (89.2%, 95% CI: 86.4–91.5; **Table S4B**), and low in both Control (9.3%, 7.1–12.2) and D1-inoculated recruits (8.1%, 6.2–10.4). C1-inoculated recruits differed significantly from both other groups (Tukey-adjusted p < 0.001), whereas Control and D1-inoculated recruits did not differ from one another (odds ratio = 1.17, p = 0.603; **Table S4D**).

**Figure 4.**
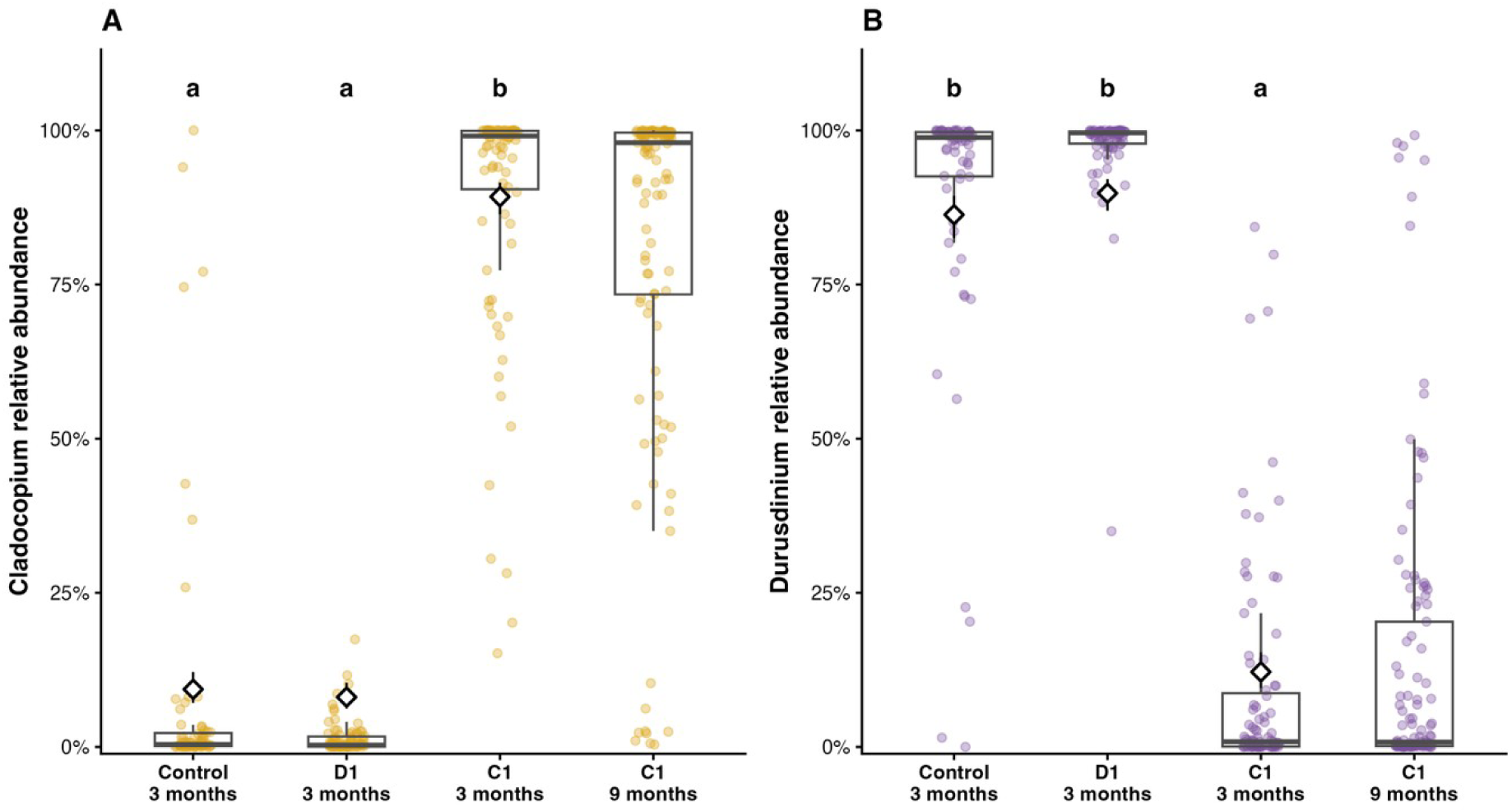
Relative abundance of (A) Cladocopium and (B) Durusdinium in recruits three and nine months after deployment. Relative abundance is the proportion of ITS2 sequence reads assigned to each genus per recruit. Small dots show individual recruits (three months, n = 249; nine months, n = 117). Recruits sampled at three and nine months are different individuals, not repeated measurements. Boxplots show the median, interquartile range and range within 1.5 × the interquartile range of the raw data. Diamonds and error bars show the model-predicted mean ± 95% CI from the beta GLMM for the three-month treatment comparison (Control, C1, D1). Letters indicate significant differences among the three-month treatments (Tukey-adjusted pairwise comparisons, p < 0.05; groups sharing a letter do not differ). C1 at nine months was analysed in a separate model comparing three and nine months within C1 (not significant for either genus: Cladocopium p = 0.072, Durusdinium p = 0.477) and is not part of the lettering.

Treatment also had a significant effect on *Durusdinium* relative abundance (Wald χ^2^_2_ = 355.73, p < 0.001; **Figure 4B; Table S4C**). Both Control (86.3%, 95% CI: 82.5–89.4) and D1-inoculated recruits (89.8%, 87.0–92.1) were strongly dominated by *Durusdinium* (odds ratio = 0.72, p = 0.123), indicating that both treatments converged on similar genus composition despite differing inoculation histories, while *Durusdinium* relative abundance in the C1 treatment remained low (12.2%, 9.6–15.3). Pairwise comparisons confirmed that C1-inoculated recruits differed strongly from both Control and D1 treatments for *Durusdinium* (Tukey-adjusted p < 0.001 in all cases; **Table S4D**), while *Durusdinium* in Control and D1 were not significantly different.

After nine months on the reef, neither *Cladocopium* nor *Durusdinium* relative abundance differed significantly between C1-inoculated recruits sampled at three and nine months (*Cladocopium*: Wald χ^2^_1_ = 3.25, odds ratio = 1.33, p = 0.072; *Durusdinium*: Wald χ^2^_1_ = 0.50, odds ratio = 0.90, p = 0.477; **Table S4G**). A small difference in the predicted means was nevertheless apparent: model-predicted *Cladocopium* relative abundance was 83.3% (95% CI: 79.1–86.8) at three months and 78.9% (74.6–82.7) at nine months, while *Durusdinium* relative abundance was 14.4% (11.3–18.2) and 15.8% (12.7–19.5; **Table S4F; Figure 4A–B**).

### Inoculated symbiont culture fingerprints persisted in field-deployed recruits

C1-inoculated recruits showed strong retention of the fingerprint at three months but with sample-to-sample heterogeneity: the divergence between mean and median fingerprint contribution (69.1% vs. 95.4%) reflected a bimodal pattern, whereby most recruits maintained near-complete inoculum dominance while a minority hosted more divergent *Cladocopium* communities (**Figure 5B; Table S5B**). Similarity to D1 fingerprint remained low but non-negligible in a small subset of samples, and low-abundance non-fingerprint taxa were detected without displacing the dominant inoculum: *Durusdinium* in most recruits (89%; median relative abundance 0.24%) and *Symbiodinium* in about half (51%; 0.09%), whereas *Fugacium* (9%; 0.08%) and *Breviolum* (1%; 0.22%) were rare (**Table S6**).

**Figure 5.**
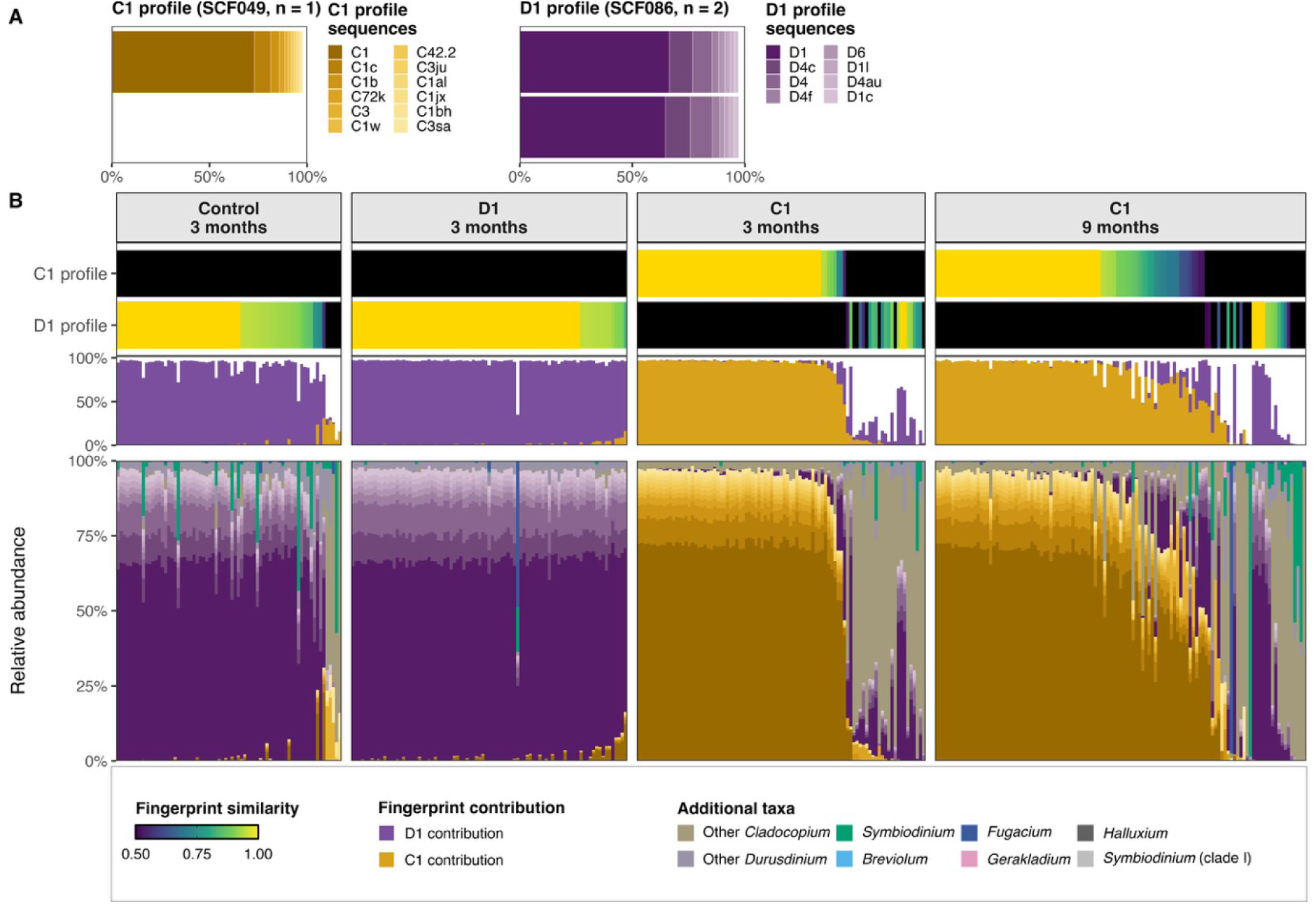
Culture-specific ITS2 fingerprint similarity and composition of recruits three and nine months after deployment. A fingerprint is the set of the most abundant ITS2 sequences that together make up 98% of the signal of an inoculated culture (12 sequences for the C1 culture SCF049, 8 for the D1 culture SCF086). (A) ITS2 sequence composition of the C1 (SCF049, n = 1 replicate) and D1 (SCF086, n = 2 replicates) culture profiles, with sequences in order of abundance (ochre shades, *Cladocopium* C1 profile; purple shades, *Durusdinium* D1 profile). (B) For each recruit, from top to bottom: fingerprint similarity to the C1 and D1 profiles (colour strip; similarity is 1 − Bray–Curtis dissimilarity; gold indicates similarity ≥ 0.95, black indicates similarity < 0.50, with intermediate values shown as a colour gradient); fingerprint contribution, the proportion of ITS2 reads matching each culture fingerprint, stacked cumulatively (C1 ochre, D1 purple); and relative ITS2 sequence composition. Recruits are ordered within each treatment and timepoint by fingerprint similarity. Total recruit samples shown, n = 366.

D1-inoculated recruits showed a more uniform pattern: mean and median fingerprint contribution were concordant (94.6% vs. 96.4%), consistent with near-uniform inoculum dominance across nearly all recruits and negligible similarity to the C1 fingerprint (**Figure 5B; Table S5B**). Low-abundance non-fingerprint taxa were also detected, in particular *Durusdinium* (all recruits; median 2.74%), *Cladocopium* (86%; 0.23%) and *Symbiodinium* (72%; 0.07%), whereas *Fugacium* was less common (17%; 0.03%); all remained at low relative abundance throughout (**Table S6).**

Control recruits showed no similarity to the C1 fingerprint (**Table S5B**). Communities were instead strongly dominated by the D1 fingerprint, though a wider divergence between mean and median D1 fingerprint contribution (84.0% vs. 94.8%) than in D1-inoculated recruits indicates Control recruits hosted more heterogeneous communities (**Table S5B**).

At nine months, the C1 fingerprint remained prevalent but recruits sampled at nine months had lower median fingerprint similarity to the SCF049 profile (0.91) than those sampled at three months (0.97), and lower median fingerprint contribution (87.3% vs. 95.4%; **Table S5B; Figure 5B**), consistent with acquisition of non-inoculum symbionts. A minority of C1 recruits showed close correspondence to the D1 fingerprint profile or approximately equal *Cladocopium– Durusdinium* co-dominance by nine months, and background taxa, including *Gerakladium* and *Halluxium*, were detected only at nine months (**Table S6**).

## DISCUSSION

### Inoculated symbionts are retained by recruits following field deployment

At the genus level, inoculation produced a strong imprinting of symbiont composition in C1-inoculated recruits, which remained *Cladocopium*-dominated among recruits sampled at three and nine months, whereas *Durusdinium* dominated both D1-inoculated and Control recruits after three months. The fingerprint approach showed that this pattern reflected a retention of the cultured ITS2 assemblage rather than uptake of a different species or lineage within the same genus. A substantial minority of C1-inoculated recruits hosted more compositionally diverse communities by three months. Because ITS2 does not resolve strains within a species, and Control recruits naturally acquired *Durusdinium* resembling D1, we interpret the symbionts taken up by the control as consistent with the inoculated strains but are unable to confirm strain-level identity.

Early colonisation by inoculum appeared to limit subsequent colonisation by naturally occurring symbionts in the C1 treatment. This is consistent with a priority effect (Fukami 2015) and supported by two lines of evidence. First, if *Durusdinium* dominance of juvenile corals is the natural background state at Davies Reef, D1 inoculation likely reinforced the outcome recruits would have reached regardless of treatment, as most D1-inoculated and Control recruits closely matched the SCF086 fingerprint, whereas C1-inoculated recruits diverged sharply from this state. While non-culture *Durusdinium* sequences were detected in most C1 recruits, it remained as a minority group within these samples up to nine months, consistent with an established *Cladocopium* population suppressing further colonisation despite ongoing exposure.

Secondly, symbionts in Control recruits may have originated from D1-inoculated recruits as Control, D1- and C1-inoculated devices were held together for one week before deployment and were deployed within the same 1 m^2^ reef plots. Control and C1-inoculated recruits therefore had similar opportunities to acquire *Durusdinium* both before and after deployment. While the C1 treatment showed traces of the D1 culture, only Control and D1 recruits became *Durusdinium*-dominated.

Pre-emption is a plausible mechanism here, as inoculation allowed the inoculated lineage to occupy host niches before competing natural symbionts arrived, as documented in other host-associated microbial systems, including early-life gut microbiota (Sprockett et al. 2018; Debray et al. 2022).

### Fingerprint analyses improved detection and interpretation of inoculum community changes

For D1-inoculated recruits, genus-level *Durusdinium* dominance alone could not resolve inoculum retention, because *Durusdinium* also dominated Controls. However, fingerprint similarity indicated that a higher proportion of D1-inoculated recruits exceeded the high-similarity threshold to cultures compared to Controls. The *Durusdinium*-dominated Controls admit two non-exclusive explanations. First, aposymbiotic recruits may preferentially acquire *Durusdinium* from a mixed symbiont pool. This is consistent with field observations that juvenile *Acropora* rapidly become *Durusdinium*-dominated regardless of the symbionts hosted by adults (Abrego et al. 2009a) and with experimental mixed-inoculum exposure (Ivory et al. 2025). Second, because SCF086 was isolated from Davies Reef, Control recruits may have acquired environmental *Durusdinium* that ITS2 cannot distinguish from the cultured strain (Hume et al. 2019; Davies et al. 2023), so the two sources cannot be separated with sequence data alone. The fingerprint nonetheless suggests that Control dominance was not compositionally uniform, with more species diversity than within D1 treatment, consistent with acquisition from the diverse environmental pool and mixed juvenile communities reported (Quigley et al. 2017; Fujise et al. 2021), although some contribution from uniform uptake of one strain cannot be excluded.

A further limitation applies to both explanations. Symbiont composition could only be characterised destructively in surviving recruits, so we cannot determine whether Control recruits that initially acquired *Cladocopium* died before sampling or were displaced by *Durusdinium*. Resolving this would require sampling seawater from larval rearing, settlement, device-assembly and grow-out tanks, and coral tissue before inoculation, immediately after inoculation and immediately before deployment, which would separate pre-deployment cross-infection, in-facility acquisition and post-deployment reef acquisition. Tracking cultured strains within a species will ultimately require higher-resolution genotyping (Davies et al. 2023). More broadly, these results come from one coral species at one reef over nine months, so testing across species, reef contexts and through to adulthood is a priority. Recruits sampled at nine months differed in symbiont composition from those sampled at three months, although this difference was not statistically detectable at the genus level. The genus-level models also fitted imperfectly, because relative abundances were strongly bimodal, so their p-values and confidence intervals are approximate.

The treatment differences were large and agreed with the fingerprint analyses, whereas the non-significant change between three and nine months in C1-inoculated recruits should be read cautiously. Fingerprint data suggest that non-fingerprint *Cladocopium* variants, possibly environmentally acquired, were already present in all C1-inoculated recruits at three months, though at low median abundance. By nine months, a small proportion of recruits closely matched the D1 fingerprint profile, indicating that some recruits hosted a community resembling D1. A further subset of recruits showed approximately equal co-dominance of *Cladocopium* and *Durusdinium*. Because sampling was destructive and the nine-month recruits are surviving individuals distinct from those sampled at three months, these differences could also reflect differential survival or differences among individual recruits rather than change within individuals. One possibility is that *Durusdinium* and non-fingerprint *Cladocopium* lineages may colonise established recruits more effectively than the cultured C1 strain, consistent with the flexible associations and changes in dominant symbionts reported during early ontogeny in *Acropora* (Little et al. 2004; Abrego et al. 2009b). Because all D1 devices were retrieved at three months, nine-month data are unavailable for D1-inoculated recruits, so we cannot test whether *Durusdinium* communities would show comparable stability.

### Symbiont inoculation improves early recruit survival

Both inoculation treatments more than doubled survival relative to Controls. Notably, D1-inoculated and Control recruits harboured statistically indistinguishable *Durusdinium*-dominated communities by three months, yet their survival differed. This suggests that the early symbiosis establishment through inoculation, is an important determinant of early survival. This is consistent with early symbiont provisioning accelerating colonisation, enhancing growth and survivorship relative to aposymbiotic juveniles (Cumbo et al. 2013), and with Symbiodiniaceae genera’s specific infectivity (Pollock et al. 2017). Studies using SCF086 and other experimentally evolved thermotolerant strains similarly show introduced Symbiodiniaceae alter juvenile fitness under controlled conditions (Nitschke et al. 2024; Ivory et al. 2025; Lamb et al. 2026). Our results extend these findings to the field, showing that inoculated symbionts remained dominant in recruits under natural conditions, including a transient temperature fluctuation shortly after deployment, and translates into substantially improved survival. To our knowledge, this is the first field study to inoculate device-deployed, sexually propagated recruits with cultured symbionts and characterise the inoculated symbionts alongside survival. The temperature record suggests one possible cause. Within a month of deployment, a brief cooling coincided with a peak in current speed, followed by a rise of ∼1.5°C. Inoculated recruits may have been better placed to buffer this fluctuation compared to Controls that were potentially still acquiring symbionts from the environment. This remains speculative, as we cannot separate it from other sources of early mortality, and symbionts were characterised only in surviving recruits. Testing it would require non-destructive symbiont detection combined with survival tracking of the same individuals over time.

A plausible mechanism is energetic. Settlement and early skeletal formation are energetically demanding, and recruits become increasingly dependent on photosymbiont-derived nutrition as maternal reserves are exhausted (Huffmyer et al. 2025). Delivering high-density cultured symbionts before deployment may have shortened this vulnerable window, allowing recruits to reach a positive energy balance earlier, with potential benefits for growth and calcification (Harii et al. 2010; Cumbo et al. 2013). Direct measurement of growth, calcification, and bioenergetics alongside inoculation is needed to test this. The thermal-tolerance advantage of *Durusdinium* is documented under sustained heat stress in adult corals (Berkelmans & van Oppen 2006; Jones & Berkelmans 2010) and in *Acropora spathulata* juveniles at 31°C (Quigley et al. 2020). The transient fluctuation recorded after deployment at Davies was brief, and C1 and D1 survival did not differ. This suggests that successful establishment of symbiosis, rather than genus-specific thermal traits, was the primary determinant of early survival under these conditions. Low-abundance background *Durusdinium* in most C1-inoculated recruits may also provide a reservoir for community shuffling towards greater thermal tolerance under stress (Cunning et al. 2015). A prolonged thermal challenge assay would test both whether a D1 advantage emerges under sustained heat and whether this reservoir allows C1-inoculated recruits to compensate. For restoration, C1 strains capable of successful early colonisation may therefore offer survival benefits at least equivalent to D1 where sustained thermal stress is not anticipated, although *Durusdinium* dominated even under mixed-inoculum exposure in Ivory et al. (2025).

Habitat context further modulated the survival benefit of inoculation. Effect sizes across the habitat gradients were large for both inoculated treatments but opposite in direction. D1-inoculated recruits showed large effects across CCA, sediment, sand, and soft coral habitats, whereas C1-inoculated recruits showed the reverse pattern, with the largest single effect for CCA. Control recruits responded little to sediment, sand, and soft coral cover. This pattern should not be read as evidence that sediment, sand, or soft coral cover benefits C1-inoculated recruits. Sedimentation and competitive overgrowth are well-documented drivers of recruit mortality (Kuffner et al. 2006; Penin et al. 2011), which makes a positive effect biologically implausible. A more plausible explanation is reduced vulnerability: larger or better-established C1 recruits may be less susceptible to burial or smothering at a given cover level (Brunner et al. 2021), without the habitat conferring any advantage. This remains speculative, because recruit size and condition were not measured.

Plot-level cover also matches poorly the scale at which these factors act. Sediment, sand, and soft coral are likely to affect recruits locally, at the scale of individual devices or tabs, and devices are mostly retained in place once deployed (Ramsby et al. 2026b), so the conditions a recruit experiences are set at the device scale. A plot-averaged predictor may therefore poorly represent the cover an individual recruit experienced. Consistent with this, device-scale variance exceeded plot-scale variance, indicating that microhabitat heterogeneity below our survey resolution was the dominant source of unexplained variation in survival, in line with Page et al. (2024) and Jurriaans et al. (2025, 2026). They both found recruit performance shaped primarily at the device-scale. These associations point to a complex, scale-limited relationship between habitat and inoculation outcome, possibly confounded by treatment-associated differences in recruit condition that were not measured here.

### Restoration implications of symbiont inoculations

Symbiont inoculation substantially improved early recruit survival, and C1 inoculation produced *Cladocopium* dominance that persisted for at least nine months. Comparable survival of C1- and D1-inoculated recruits suggests that the timing and success of symbiont establishment can matter more than genus identity, while the modulation of benefits by benthic context shows that restoration outcomes depend on both symbiont provisioning and fine-scale deployment environment. The fingerprint framework developed here provides a quantitative tool for assessing inoculum retention in future field studies, although tracking cultured strains within a species will ultimately require higher-resolution genotyping (Davies et al. 2023). These results come from one coral species at one reef over nine months, so testing across species, reef contexts and through adulthood is a priority.

## ACKNOWLEDGEMENTS

We acknowledge the Bindal and Wulgurukaba as the Traditional Custodians of the land and sea Countries where this research took place and pay our respects to elders past and present. We thank them for providing Free Prior Informed Consent to undertake this work. Coral collections, deployments and experiments were conducted under GBRMPA permit G49457.1. Funding was provided by the Reef Restoration and Adaptation Program (RRAP), which was funded by a partnership between the Australian Government’s Reef Trust and the Great Barrier Reef Foundation. We thank Elizabeth Ivory for her laboratory support, and the vessel crews, SeaSim and AIMS staff, students, collaborators and volunteers who supported various aspects of the RRAP Coral Aquaculture and Deployment research sub-program.

## Data availability

ITS2 sequences FASTQ will be deposited in the NCBI Sequence Read Archive, SymPortal output and R script will be available on GitHub upon acceptance.

## Conflict of interest

The authors declare no conflicts of interest.

## Author contributions

MAW, MRN, YS conceived and designed the research; SF, EP performed the experiments; EP, MRN analysed the data; EP wrote the manuscript; all authors edited the manuscript.

## Notes

### Competing Interest Statement

The authors have declared no competing interest.

## REFERENCES

Abrego D, van Oppen MJH, Willis BL (2009a) Highly infectious symbiont dominates initial uptake in coral juveniles. Molecular Ecology 18:3518–3531.

Abrego D, van Oppen MJH, Willis BL (2009b) Onset of algal endosymbiont specificity varies among closely related species of *Acropora* corals during early ontogeny. Molecular Ecology 18:3532–3543.

Abrego D, Willis BL, van Oppen MJH (2012) Impact of light and temperature on the uptake of algal symbionts by coral juveniles. PLoS ONE 7:e50311.

Australian Institute of Marine Science (2024) ReefCloud. 10.25845/g5gk-ty57

Baird AH, Guest JR, Willis BL (2009) Systematic and biogeographical patterns in the reproductive biology of scleractinian corals. Annual Review of Ecology, Evolution, and Systematics 40:551–571.

Baker AC (2003) Flexibility and specificity in coral–algal symbiosis: diversity, ecology, and biogeography of *Symbiodinium*. Annual Review of Ecology, Evolution, and Systematics 34:661–689.

Bayraktarov E, Saunders MI, Abdullah S, Mills M, Beher J, Possingham HP, Mumby PJ, Lovelock CE (2016) The cost and feasibility of marine coastal restoration. Ecological Applications 26:1055–1074.

Berkelmans R, van Oppen MJH (2006) The role of zooxanthellae in the thermal tolerance of corals: a ‘nugget of hope’ for coral reefs in an era of climate change. Proceedings of the Royal Society B: Biological Sciences 273:2305–2312.

Boström-Einarsson L, Babcock RC, Bayraktarov E, Ceccarelli D, Cook N, Ferse SCA, Hancock B, Harrison P, Hein M, Shaver E, Smith A, Suggett D, Stewart-Sinclair PJ, Vardi T, McLeod IM (2020) Coral restoration – a systematic review of current methods, successes, failures and future directions. PLOS ONE 15:e0226631.

Brunner CA, Uthicke S, Ricardo GF, Hoogenboom MO, Negri AP (2021) Climate change doubles sedimentation-induced coral recruit mortality. Science of the Total Environment 768:143897.

Chamberland VF, Petersen D, Guest JR, Petersen U, Brittsan M, Vermeij MJA (2017) New seeding approach reduces costs and time to outplant sexually propagated corals for reef restoration. Scientific Reports 7:18076.

Cumbo VR, Baird AH, van Oppen MJH (2013) The promiscuous larvae: flexibility in the establishment of symbiosis in corals. Coral Reefs 32:111–120.

Cunning R, Silverstein RN, Baker AC (2015) Investigating the causes and consequences of symbiont shuffling in a multi-partner reef coral symbiosis under environmental change. Proceedings of the Royal Society B: Biological Sciences 282:20141725.

Dallmeyer-Drennen G, Sato Y, Page CA, Bourne DG, Smith HA (2026) Genetic diversity of sexually propagated corals is maintained from the aquarium to the reef. Molecular Ecology 35:e70289.

Davies SW, Gamache MH, Howe-Kerr LI, Kriefall NG, Baker AC, Banaszak AT, Bay LK, Bellantuono AJ, Bhattacharya D, Chan CX, Claar DC, Coffroth MA, Cunning R, Davy SK, del Campo J, Díaz-Almeyda EM, Frommlet JC, Fuess LE, González-Pech RA, Goulet TL, Hoadley KD, Howells EJ, Hume BCC, Kemp DW, Kenkel CD, Kitchen SA, LaJeunesse TC, Lin S, McIlroy SE, McMinds R, Nitschke MR, Oakley CA, Peixoto RS, Prada C, Putnam HM, Quigley K, Reich HG, Reimer JD, Rodriguez-Lanetty M, Rosales SM, Saad OS, Sampayo EM, Santos SR, Shoguchi E, Smith EG, Stat M, Stephens TG, Strader ME, Suggett DJ, Swain TD, Tran C, Traylor-Knowles N, Voolstra CR, Warner ME, Weis VM, Wright RM, Xiang T, Yamashita H, Ziegler M, Correa AMS, Parkinson JE (2023) Building consensus around the assessment and interpretation of Symbiodiniaceae diversity. PeerJ 11:e15023.

Debray R, Herbert RA, Jaffe AL, Crits-Christoph A, Power ME, Koskella B (2022) Priority effects in microbiome assembly. Nature Reviews Microbiology 20:109–121.

Doropoulos C, Vons F, Elzinga J, ter Hofstede R, Salee K, van Koningsveld M, Babcock RC (2019a) Testing industrial-scale coral restoration techniques: harvesting and culturing wild coral-spawn slicks. Frontiers in Marine Science 6.

Doropoulos C, Elzinga J, ter Hofstede R, van Koningsveld M, Babcock RC (2019b) Optimizing industrial-scale coral reef restoration: comparing harvesting wild coral spawn slicks and transplanting gravid adult colonies. Restoration Ecology 27:758–767.

Fujise L, Suggett DJ, Stat M, Kahlke T, Bunce M, Gardner SG, Goyen S, Woodcock S, Ralph PJ, Seymour JR, Siboni N, Nitschke MR (2021) Unlocking the phylogenetic diversity, primary habitats, and abundances of free-living Symbiodiniaceae on a coral reef. Molecular Ecology 30:343–360.

Fukami T (2015) Historical contingency in community assembly: integrating niches, species pools, and priority effects. Annual Review of Ecology, Evolution, and Systematics 46:1–23.

Harii S, Yamamoto M, Hoegh-Guldberg O (2010) The relative contribution of dinoflagellate photosynthesis and stored lipids to the survivorship of symbiotic larvae of the reef-building corals. Marine Biology 157:1215–1224.

Huffmyer AS, Wong KH, Becker DM, Strand E, Mass T, Putnam HM (2025) Shifts and critical periods in coral metabolism reveal energetic vulnerability during development. Current Biology 35:2858–2871.e6.

Hughes TP, Kerry JT, Baird AH, Connolly SR, Dietzel A, Eakin CM, Heron SF, Hoey AS, Hoogenboom MO, Liu G, McWilliam MJ, Pears RJ, Pratchett MS, Skirving WJ, Stella JS, Torda G (2018) Global warming transforms coral reef assemblages. Nature 556:492–496.

Humanes A, Fabricius KE, Ferrari R, Ortiz JC (2025) Ecological quantitative criteria for reef site prioritisation to maximise survivorship and growth of outplanted corals. Journal of Environmental Management 392:126585.

Hume BCC, Ziegler M, Poulain J, Pochon X, Romac S, Boissin E, de Vargas C, Planes S, Wincker P, Voolstra CR (2018) An improved primer set and amplification protocol with increased specificity and sensitivity targeting the *Symbiodinium* ITS2 region. PeerJ 6:e4816.

Hume BCC, Smith EG, Ziegler M, Warrington HJM, Burt JA, LaJeunesse TC, Wiedenmann J, Voolstra CR (2019) SymPortal: a novel analytical framework and platform for coral algal symbiont next-generation sequencing ITS2 profiling. Molecular Ecology Resources 19:1063–1080.

Ivory EA, Mieog JC, Nitschke MR, van Oppen MJH, Abrego D (2025) Interactions among wildtype and heat-evolved photosymbionts shape performance of coral recruits. Coral Reefs 44:643–655.

Jones A, Berkelmans R (2010) Potential costs of acclimatization to a warmer climate: growth of a reef coral with heat tolerant vs. sensitive symbiont types. PLoS ONE 5:e10437.

Jurriaans S, Lefèvre CD, Allen K, Giuliano C, Page CA, Puotinen M, Radford B, Sims CA, Whitman TN, Randall CJ (2025) Wave energy and other environmental drivers as predictors of seeded-coral performance on the Great Barrier Reef. Scientific Reports 15:38335.

Jurriaans S, Lefèvre CD, Ferguson S, Randall CJ (2026) Propagation method and species drive survival patterns across reef zones in coral seeding on the Great Barrier Reef. Restoration Ecology 34:e70358.

Kuffner I, Walters L, Becerro M, Paul V, Ritson-Williams R, Beach K (2006) Inhibition of coral recruitment by macroalgae and cyanobacteria. Marine Ecology Progress Series 323:107– 117.

LaJeunesse TC, Parkinson JE, Gabrielson PW, Jeong HJ, Reimer JD, Voolstra CR, Santos SR (2018) Systematic revision of Symbiodiniaceae highlights the antiquity and diversity of coral endosymbionts. Current Biology 28:2570–2580.e6.

Lamb AM, Morgans CA, McCutchan GA, Mocellin VJL, Denis H, Naugle MS, Lemonius AJ, Edmunds RC, Wooster MK, Bay LK, Howells EJ, van Oppen MJH (2026) A selective breeding design based on parental rapid heat stress thresholds did not produce more heat-tolerant coral larvae. Ecological Solutions and Evidence 7:e70178.

Little AF, van Oppen MJH, Willis BL (2004) Flexibility in algal endosymbioses shapes growth in reef corals. Science 304:1492–1494.

Mendoza Quiroz S, Banaszak AT, McGonigle ML, Bickel AR, Guest JR, Miller MW (2025) Influence of attachment techniques on coral seeding unit deployment cost and performance. Restoration Ecology 33:e70090.

Muscatine L, Porter JW (1977) Reef corals: mutualistic symbioses adapted to nutrient-poor environments. BioScience 27:454–460.

Nitschke MR, Abrego D, Allen CE, Alvarez-Roa C, Boulotte NM, Buerger P, Chan WY, Fae Neto WA, Ivory E, Johnston B, Meyers L, Parra VC, Peplow L, Perez T, Scharfenstein HJ, van Oppen MJH (2024) The use of experimentally evolved coral photosymbionts for reef restoration. Trends in Microbiology 32:1241–1252.

Page CA, Giuliano C, Randall CJ (2024) Benthic communities influence coral seeding success at fine spatial scales. Restoration Ecology 32:e14212.

Penin L, Michonneau F, Baird A, Connolly S, Pratchett M, Kayal M, Adjeroud M (2010) Early post-settlement mortality and the structure of coral assemblages. Marine Ecology Progress Series 408:55–64.

Penin L, Michonneau F, Carroll A, Adjeroud M (2011) Effects of predators and grazers exclusion on early post-settlement coral mortality. Hydrobiologia 663:259–264.

Pollock FJ, Katz SM, van de Water JAJM, Davies SW, Hein M, Torda G, Matz MV, Beltran VH, Buerger P, Puill-Stephan E, Abrego D, Bourne DG, Willis BL (2017) Coral larvae for restoration and research: a large-scale method for rearing *Acropora millepora* larvae, inducing settlement, and establishing symbiosis. PeerJ 5:e3732.

Quigley KM, Baker AC, Coffroth MA, Willis BL, van Oppen MJH (2018) Bleaching resistance and the role of algal endosymbionts. Pages 111–151. In: van Oppen MJH, Lough JM (eds) Coral bleaching: patterns, processes, causes and consequences. Springer International Publishing, Cham.

Quigley KM, Randall CJ, van Oppen MJH, Bay LK (2020) Assessing the role of historical temperature regime and algal symbionts on the heat tolerance of coral juveniles. Biology Open 9:bio047316.

Quigley KM, Willis BL, Bay LK (2017) Heritability of the *Symbiodinium* community in vertically- and horizontally-transmitting broadcast spawning corals. Scientific Reports 7:8219.

R Core Team (2026) R: a language and environment for statistical computing. R Foundation for Statistical Computing, Vienna, Austria. https://www.R-project.org/

Ramsby BD, Brunner R, Barton J, Ferguson S, Grimm C, Hiçyilmaz YC, Bourne DG, Negri AP, Severati A, Sato Y, Abdul Wahab MA (2026a) Coral larval aquaculture: species-specific survival and microbial dynamics in flow-through systems. PLOS ONE 21:e0340422.

Ramsby BD, Forster R, Ferguson SN, Haikola P, Randall CJ, Abdul Wahab MA, Mead DJ, Severati A (2026b) Developing coral seeding devices and rapid deployment methods to scale up reef restoration. Restoration Ecology 34:e70206.

Severati A, Nordborg FM, Heyward A, Abdul Wahab MA, Brunner CA, Montalvo-Proano J, Negri AP (2024) The AutoSpawner system – automated *ex situ* spawning and fertilisation of corals for reef restoration. Journal of Environmental Management 366:121886.

Sprockett D, Fukami T, Relman DA (2018) Role of priority effects in the early-life assembly of the gut microbiota. Nature Reviews Gastroenterology & Hepatology 15:197–205.

Vermeij MJA, Sandin SA (2008) Density-dependent settlement and mortality structure the earliest life phases of a coral population. Ecology 89:1994–2004.

Waters C, Harrison PL, Gouezo M, Severati A, Doropoulos C (2025) Early-stage coral survivorship using wild larval assemblages on coral seeding devices for reef restoration. Restoration Ecology 33:e14387.

Whitman TN, Jurriaans S, Lefevre C, Sims CA, Radford B, Puotinen M, Hoogenboom MO, Negri AP, Randall CJ (2025) Seeded *Acropora digitifera* corals survive best on wave-exposed reefs with grazing from small fishes. Restoration Ecology 33:e70016.

Yellowlees D, Rees TAV, Leggat W (2008) Metabolic interactions between algal symbionts and invertebrate hosts. Plant, Cell & Environment 31:679–694.

