## Supplement document for "The use of cultured symbionts enhances the survival of reef-deployed coral recruits"

**Figure S1.** Daily mean (black line) ± SD (grey band) of temperature and current speed at Davies Reef, January–April 2024. Only days with complete logger records are shown.

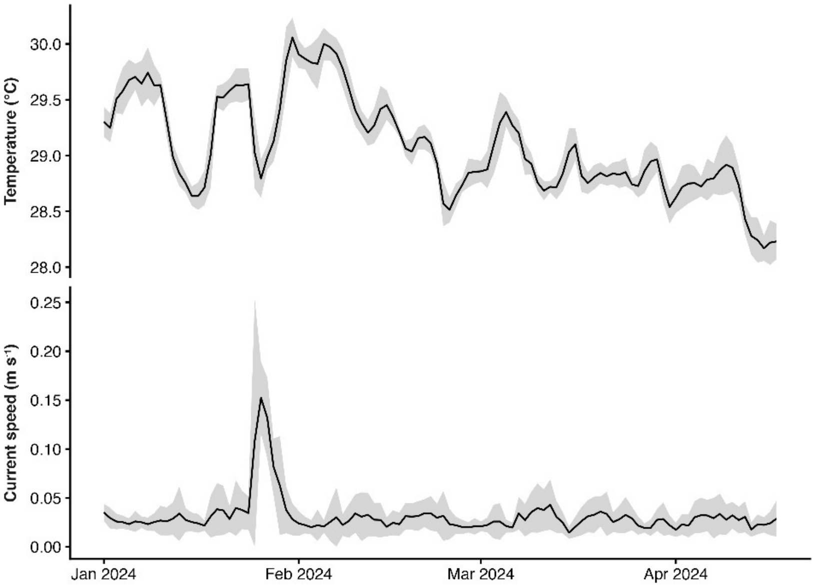

**Table S1.** Recruit observed yield and binomial GLMM (logit link) modelling of tab-level survival after three months on the reef. C1 nine-month values are descriptive only and were not included in the treatment-effect models. Control, uninoculated recruits; C1, recruits inoculated with *Cladocopium* sp. (strain SCF049); D1, recruits inoculated with *Durusdinium trenchii* (strain SCF086). GLMM, generalised linear mixed model; AIC, Akaike information criterion; SE, standard error; CI, confidence interval. Residual diagnostics for this model are shown in the "Treatment model" panels of Figure S2 and the first row of Table S2C.

**S1A.** Observed yield after three months. Level indicates whether survival was scored per device or per tab. Values are means across plots; SE is the standard error of the mean across plots.

| Level | Treatment | n plots | Mean survival (%) | SE (%) | Range (%) |
|---|---|---|---|---|---|
| Device | Control | 15 | 57.30 | 8.70 | 0.00–100.00 |
| Device | C1 | 14 | 90.50 | 4.20 | 66.70–100.00 |
| Device | D1 | 14 | 79.50 | 5.80 | 33.30–100.00 |
| Tab | Control | 15 | 31.00 | 5.40 | 0.00–61.10 |
| Tab | C1 | 14 | 63.50 | 4.40 | 44.40–88.90 |
| Tab | D1 | 14 | 57.10 | 7.10 | 11.10–94.40 |

**S1B.** Treatment effect on tab-level yield (Type II Wald $\chi^2$ test).

| Effect | n tabs | n devices | n plots | AIC | $\chi^2$ | df | p |
|---|---|---|---|---|---|---|---|
| Treatment | 630 | 210 | 15 | 788.10 | 31.99 | 2 | **<0.001** |

**S1C. Model-predicted yield (estimated marginal means from the binomial GLMM).**

| Treatment | Predicted tab-level yield (%) | SE (%) | 95% CI (%) |
|---|---|---|---|
| Control | 26.30 | 5.25 | 17.30–37.70 |
| C1 | 67.20 | 7.09 | 52.20–79.40 |
| D1 | 60.00 | 6.27 | 47.40–71.50 |

**S1D.** Pairwise contrasts, Tukey corrected.

| Contrast | Odds ratio | SE | z | p |
|---|---|---|---|---|
| Control / C1 | 0.17 | 0.06 | −4.92 | **<0.001** |
| Control / D1 | 0.24 | 0.07 | −4.80 | **<0.001** |

| Contrast | Odds ratio | SE | z | p |
|---|---|---|---|---|
| C1 / D1 | 1.36 | 0.46 | 0.92 | 0.630 |

**S1E.** Random effects.

| Random effect | Variance | SD |
|---|---|---|
| Plot | 0.42 | 0.65 |
| Device within plot | 1.26 | 1.12 |

**Table S2.** Habitat-specific binomial model testing the relationship between benthic habitat cover and tab-level recruit survival, and its interaction with treatment, fitted as fixed effects. Habitat variables were standardised (z-scored) prior to analysis. EAM – epilithic algal matrix, CCA – crustose coralline algae. Residual diagnostics for the treatment × habitat models are in Figure S2 (the five habitats with significant interactions) and Table S2C (all habitats).

**S2A.** Raw habitat cover per plot, summary statistics.

| Habitat | N plots | Min cover (%) | Max cover (%) | Mean cover (%) | SD cover (%) |
|---|---|---|---|---|---|
| EAM | 15 | 57.85 | 86.96 | 72.83 | 7.84 |
| CCA | 15 | 1.38 | 33.06 | 10.40 | 7.79 |
| Hard coral | 15 | 1.38 | 14.65 | 5.65 | 3.20 |
| Sediment | 15 | 0.83 | 13.10 | 4.46 | 3.71 |
| Sand | 15 | 0.00 | 7.53 | 1.94 | 2.56 |
| Sponges | 15 | 0.00 | 3.36 | 1.48 | 1.21 |
| Cyanobacteria | 15 | 0.00 | 3.36 | 1.07 | 1.25 |
| Other organisms | 15 | 0.00 | 5.43 | 0.85 | 1.33 |
| Macroalgae | 15 | 0.00 | 4.08 | 0.79 | 1.25 |
| Soft coral | 15 | 0.00 | 3.42 | 0.52 | 0.97 |

**S2B.** Habitat-specific models fitted for each habitat predictor. The effect size represents the model-predicted change in recruit yield (percentage points) across the observed range of habitat cover for each treatment (trt). EAM – epilithic algal matrix, CCA – crustose coralline algae. n dev., number of devices; Trt × hab., treatment × habitat interaction; Disp. p, p-value of the dispersion test.

| Habitat | n tabs | n dev. | n plots | AIC | Habitat $\chi^2$ | Habitat p | Trt × hab. $\chi^2$ | Trt × hab. p | Disp. p | Effect size Control | Effect size C1 | Effect size D1 |
|---|---|---|---|---|---|---|---|---|---|---|---|---|
| Sediment | 630 | 210 | 15 | 782.80 | 0.11 | 0.736 | 10.30 | **0.006** | 0.472 | 0.70 | 39.60 | 33.50 |
| Soft coral | 630 | 210 | 15 | 782.50 | 1.90 | 0.168 | 8.74 | **0.013** | 0.336 | 6.20 | 11.80 | 44.20 |
| CCA | 630 | 210 | 15 | 784.30 | 0.24 | 0.621 | 8.62 | **0.013** | 0.456 | 23.90 | 57.80 | 43.00 |
| Sand | 630 | 210 | 15 | 784.90 | 0.20 | 0.652 | 8.61 | **0.013** | 0.376 | 7.50 | 20.00 | 39.90 |
| Cyanobacteria | 630 | 210 | 15 | 786.70 | 1.01 | 0.314 | 6.12 | **0.047** | 0.408 | 12.40 | 19.90 | 23.20 |
| Other organisms | 630 | 210 | 15 | 785.90 | 0.31 | 0.577 | 5.92 | 0.052 | 0.424 | – | – | – |
| EAM | 630 | 210 | 15 | 788.70 | 0.01 | 0.910 | 5.22 | 0.073 | 0.448 | – | – | – |
| Macroalgae | 630 | 210 | 15 | 789.90 | 0.32 | 0.570 | 3.73 | 0.155 | 0.392 | – | – | – |
| Hard coral | 630 | 210 | 15 | 790.90 | 0.62 | 0.432 | 2.49 | 0.289 | 0.456 | – | – | – |
| Sponges | 630 | 210 | 15 | 788.60 | 3.64 | 0.056 | 2.05 | 0.359 | 0.408 | – | – | – |

**S2C.** DHARMa residual tests for all survival models. KS, Kolmogorov–Smirnov test of uniformity of scaled residuals; dispersion, nonparametric dispersion test; outliers, outlier test. All $p > 0.05$, so no departures from model assumptions were detected.

| Model | KS uniformity p | Dispersion p | Outlier p |
|---|---|---|---|
| Treatment model | 0.614 | 0.476 | 0.642 |
| Treatment × CCA | 0.698 | 0.434 | 0.642 |
| Treatment × Macroalgae | 0.562 | 0.452 | 0.642 |
| Treatment × Sand | 0.729 | 0.418 | 0.642 |
| Treatment × Sediment | 0.663 | 0.47 | 0.642 |
| Treatment × Other organisms | 0.71 | 0.458 | 0.642 |
| Treatment × Soft coral | 0.753 | 0.402 | 0.642 |
| Treatment × Hard coral | 0.579 | 0.472 | 0.642 |
| Treatment × EAM | 0.649 | 0.434 | 0.642 |
| Treatment × Sponges | 0.665 | 0.464 | 0.642 |
| Treatment × Cyanobacteria | 0.815 | 0.448 | 0.642 |

**Figure S2.** Residual diagnostics for the binomial GLMMs of survival (tab-level yield). Simulation-based scaled residuals (DHARMa; 1,000 simulations) are shown as QQ plots (left) and against model-predicted values (right) for the treatment model and for the five treatment × habitat models presented in Figure 3. Points following the diagonal in the QQ plots, and residuals scattered evenly around 0.5 without systematic pattern in the right panels, indicate that the model-simulated and observed data are consistent. No model showed significant deviation from uniformity, dispersion or outliers.

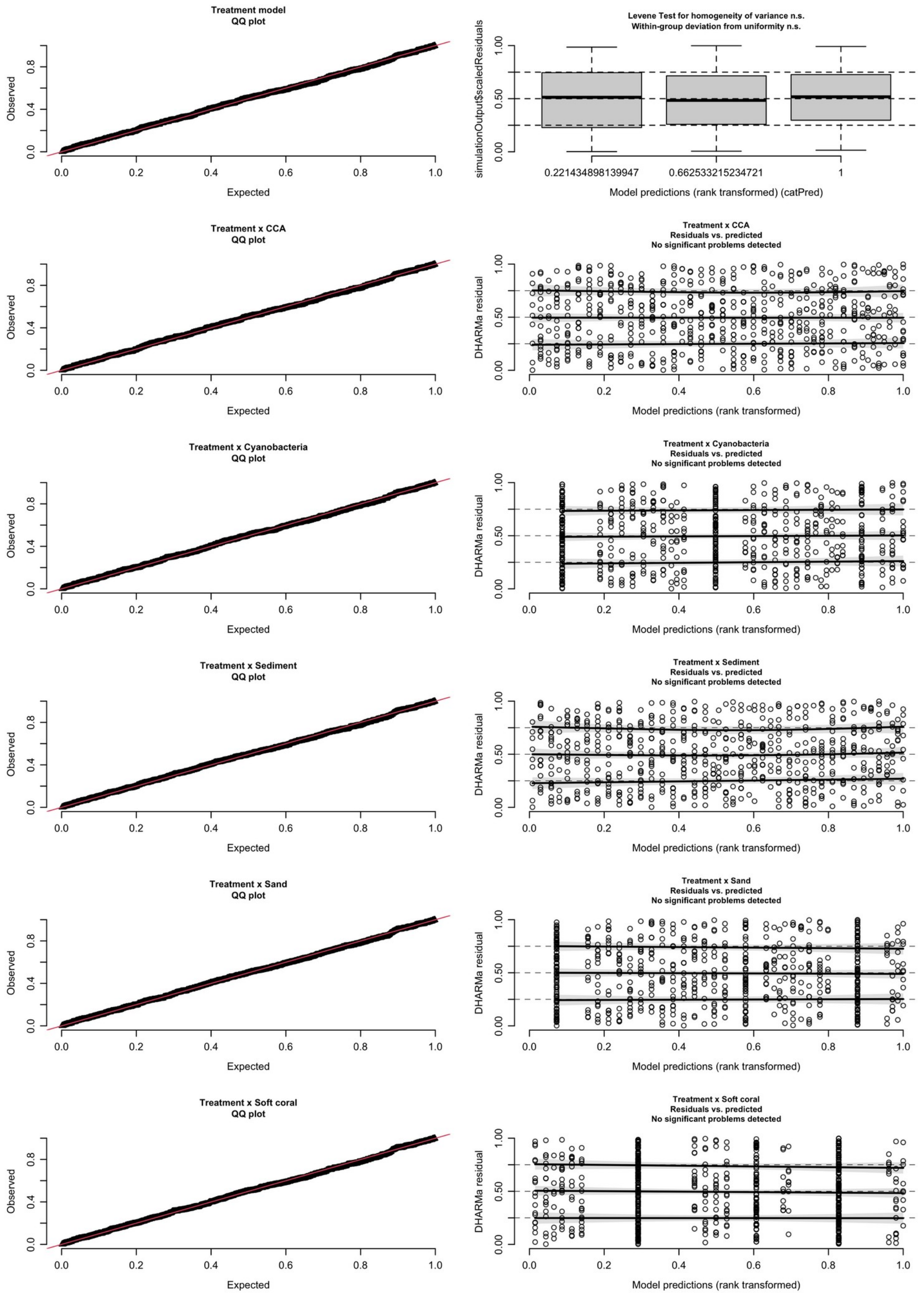

**Table S3.** Summary of successfully extracted, sequenced, and analysed recruit samples, following quality filtering. Values in parentheses indicate the initial number of samples submitted for sequencing. A value of zero indicates that samples were either not extracted, not submitted for sequencing, or did not pass quality-control filtering; it does not indicate a link to field mortality.

| Plot ID | Control (3 months) | D1 (3 months) | C1 (3 months) | C1 (9 months) |
|---|---|---|---|---|
| 235 | 12 (14) | 19 (20) | 10 (10) | 17 (21) |

| Plot ID | Control (3 months) | D1 (3 months) | C1 (3 months) | C1 (9 months) |
|---|---|---|---|---|
| 236 | 19 (19) | 11 (13) | 7 (8) | 14 (19) |
| 237 | 0 (0) | 27 (27) | 0 (0) | 11 (12) |
| 238 | 11 (11) | 0 (0) | 6 (8) | 12 (13) |
| 239 | 1 (5) | 17 (21) | 14 (15) | 5 (5) |
| 240 | 9 (13) | 13 (14) | 4 (5) | 16 (16) |
| 241 | 0 (2) | 0 (0) | 3 (3) | 6 (7) |
| 242 | 1 (4) | 0 (0) | 6 (7) | 6 (9) |
| 243 | 1 (1) | 0 (0) | 15 (15) | 4 (4) |
| 244 | 2 (3) | 0 (0) | 16 (15) | 14 (17) |
| 245 | 3 (3) | 0 (0) | 1 (1) | 0 (0) |
| 247 | 4 (4) | 0 (0) | 3 (3) | 3 (3) |
| 248 | 6 (6) | 0 (0) | 4 (4) | 4 (4) |
| 249 | 2 (9) | 0 (0) | 2 (3) | 5 (7) |
| **Total** | **71 (94)** | **87 (95)** | **91 (99)** | **117 (129)** |

**Figure S3.** Mean Symbiodiniaceae genus composition by treatment and timepoint. Bars show mean relative abundance of each genus.

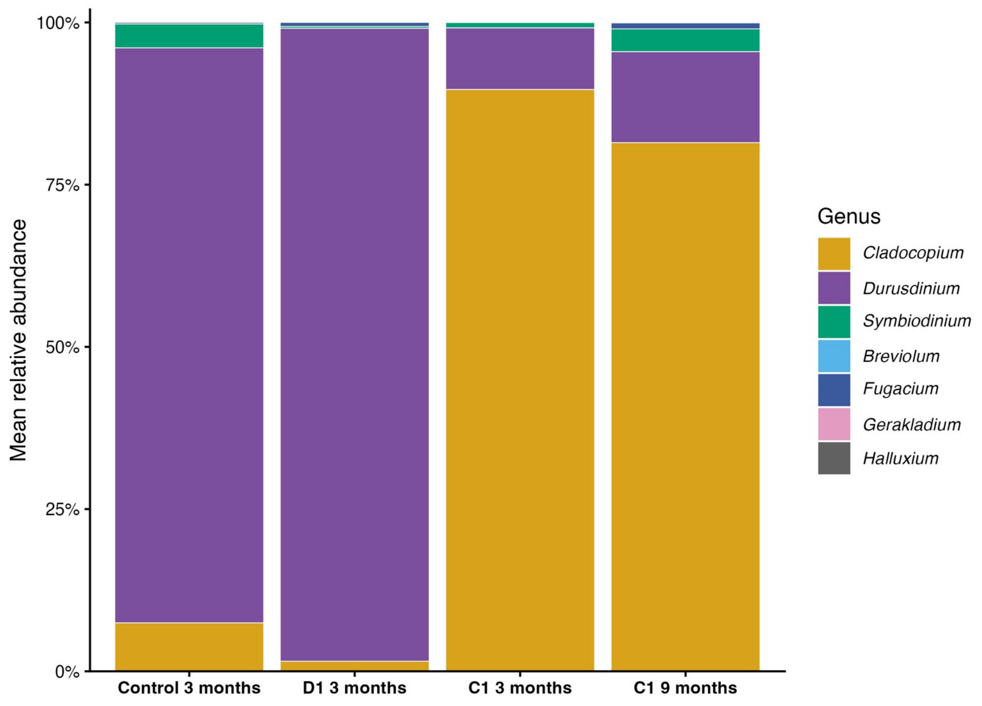

**Table S4.** Genus-level Symbiodiniaceae composition. Residual diagnostics are in Figure S4 and Table S4H.

**S4A.** Summary of genus assignments from SymPortal. Number and percentage of samples in which each Symbiodiniaceae genus was detected, out of the total number of samples per treatment and timepoint. See **Figure S2** for the corresponding mean relative abundance of each genus.

| **Genus** | **Control (3 months)** | **D1 (3 months)** | **C1 (3 months)** | **C1 (9 months)** |
|---|---|---|---|---|
| *Durusdinium* | 69/71 (97.20%) | 87/87 (100.00%) | 83/91 (91.20%) | 105/117 (89.70%) |
| *Cladocopium* | 64/71 (90.10%) | 79/87 (90.80%) | 91/91 (100.00%) | 117/117 (100.00%) |
| *Symbiodinium* | 48/71 (67.60%) | 63/87 (72.40%) | 46/91 (50.50%) | 95/117 (81.20%) |
| *Fugacium* | 26/71 (36.60%) | 15/87 (17.20%) | 8/91 (8.80%) | 35/117 (29.90%) |
| *Breviolum* | 0 | 0 | 1/91 (1.10%) | 0 |
| *Gerakladium* | 0 | 0 | 0 | 12/117 (10.30%) |
| *Halluxium* | 0 | 0 | 0 | 1/117 (0.90%) |

**S4B.** Model-predicted genus proportions for the three-month treatment comparison.

| Genus | Treatment | Estimated relative abundance % (95% CI) |
|---|---|---|
| *Cladocopium* | Control | 9.30 (7.10–12.20) |
| *Cladocopium* | C1 | 89.20 (86.40–91.50) |
| *Cladocopium* | D1 | 8.10 (6.20–10.40) |
| *Durusdinium* | Control | 86.30 (82.50–89.40) |
| *Durusdinium* | C1 | 12.20 (9.60–15.30) |
| *Durusdinium* | D1 | 89.80 (87.00–92.10) |

**S4C.** Type II Wald tests.

| Genus | $\chi^2$ | df | p |
|---|---|---|---|
| *Cladocopium* | 422.69 | 2 | **<0.001** |
| *Durusdinium* | 355.73 | 2 | **<0.001** |

**S4D.** Pairwise contrasts (Tukey adjustment).

| Genus | Contrast | Odds ratio | SE | z | p |
|---|---|---|---|---|---|
| *Cladocopium* | Control / C1 | 0.012 | 0.003 | −18.68 | **<0.001** |
| *Cladocopium* | Control / D1 | 1.173 | 0.196 | 0.96 | 0.603 |
| *Cladocopium* | C1 / D1 | 94.377 | 21.856 | 19.64 | **<0.001** |
| *Durusdinium* | Control / C1 | 45.553 | 10.456 | 16.64 | **<0.001** |
| *Durusdinium* | Control / D1 | 0.716 | 0.122 | −1.96 | 0.123 |
| *Durusdinium* | C1 / D1 | 0.016 | 0.004 | −18.16 | **<0.001** |

**S4E.** Fixed-effect coefficients on logit scale.

| Genus | Term | Estimate | SE | z | p |
|---|---|---|---|---|---|
| *Cladocopium* | (Intercept) | −2.27 | 0.15 | −15.09 | **<0.001** |
| *Cladocopium* | C1 | 4.39 | 0.24 | 18.68 | **<0.001** |
| *Cladocopium* | D1 | −0.16 | 0.17 | −0.96 | 0.338 |
| *Durusdinium* | (Intercept) | 1.84 | 0.15 | 12.46 | **<0.001** |
| *Durusdinium* | C1 | −3.82 | 0.23 | −16.64 | **<0.001** |
| *Durusdinium* | D1 | 0.33 | 0.17 | 1.96 | 0.050 |

**C1 temporal comparison (3 vs 9 months)**

The following results describe change over time within C1-inoculated recruits only, from a separate model with a different reference level to the treatment comparison above (Tables S4B–E); they should not be compared directly to it.

**S4F.** C1 temporal model: predicted genus proportions.

| Genus | Timepoint | Estimated relative abundance % (95% CI) |
|---|---|---|
| *Cladocopium* | 3 months | 83.30 (79.10–86.80) |
| *Cladocopium* | 9 months | 78.90 (74.60–82.70) |
| *Durusdinium* | 3 months | 14.40 (11.30–18.20) |
| *Durusdinium* | 9 months | 15.80 (12.70–19.50) |

**S4G.** Test of change between 3 and 9 months (no adjustment: single planned contrast).

| Genus | Odds ratio | SE | z | df | p |
|---|---|---|---|---|---|
| *Cladocopium* | 1.33 | 0.21 | 1.80 | 1 | 0.072 |
| *Durusdinium* | 0.90 | 0.14 | −0.71 | 1 | 0.477 |

**S4H.** DHARMa tests of residuals for the genus beta GLMMs: KS test of uniformity, dispersion test, outlier test and Levene's test of homogeneity of variance among model-predicted groups (p-values).

| Model | KS uniformity p | Dispersion p | Outlier p | Levene p |
|---|---|---|---|---|
| Cladocopium, treatment (3 months) | <0.001 | 0.244 | 0.089 | **<0.001** |
| Durusdinium, treatment (3 months) | <0.001 | 0.978 | 0.014 | **<0.001** |
| Cladocopium, C1 3 vs 9 months | <0.001 | 0.836 | 1 | **0.046** |
| Durusdinium, C1 3 vs 9 months | <0.001 | 0.826 | 0.066 | 0.106 |

**Figure S4**. Residual diagnostics for the beta GLMMs of genus relative abundance. Simulation-based scaled residuals (DHARMa; 1,000 simulations) are shown as QQ plots (left) and against model-predicted group (right) for *Cladocopium* and *Durusdinium* in the three-month treatment comparison and the C1 three- versus nine-month comparison. Residuals deviated from uniformity in all four models and had unequal variance among groups in three of them (Levene's test; Table S4H), consistent with the bimodal distribution of relative abundance among recruits (most near 0% or 100% of a genus, a minority intermediate). Dispersion did not differ from expectation in any model, and the outlier test was significant only for the

Durusdinium treatment model (p = 0.014). Test results are in Table S4H.

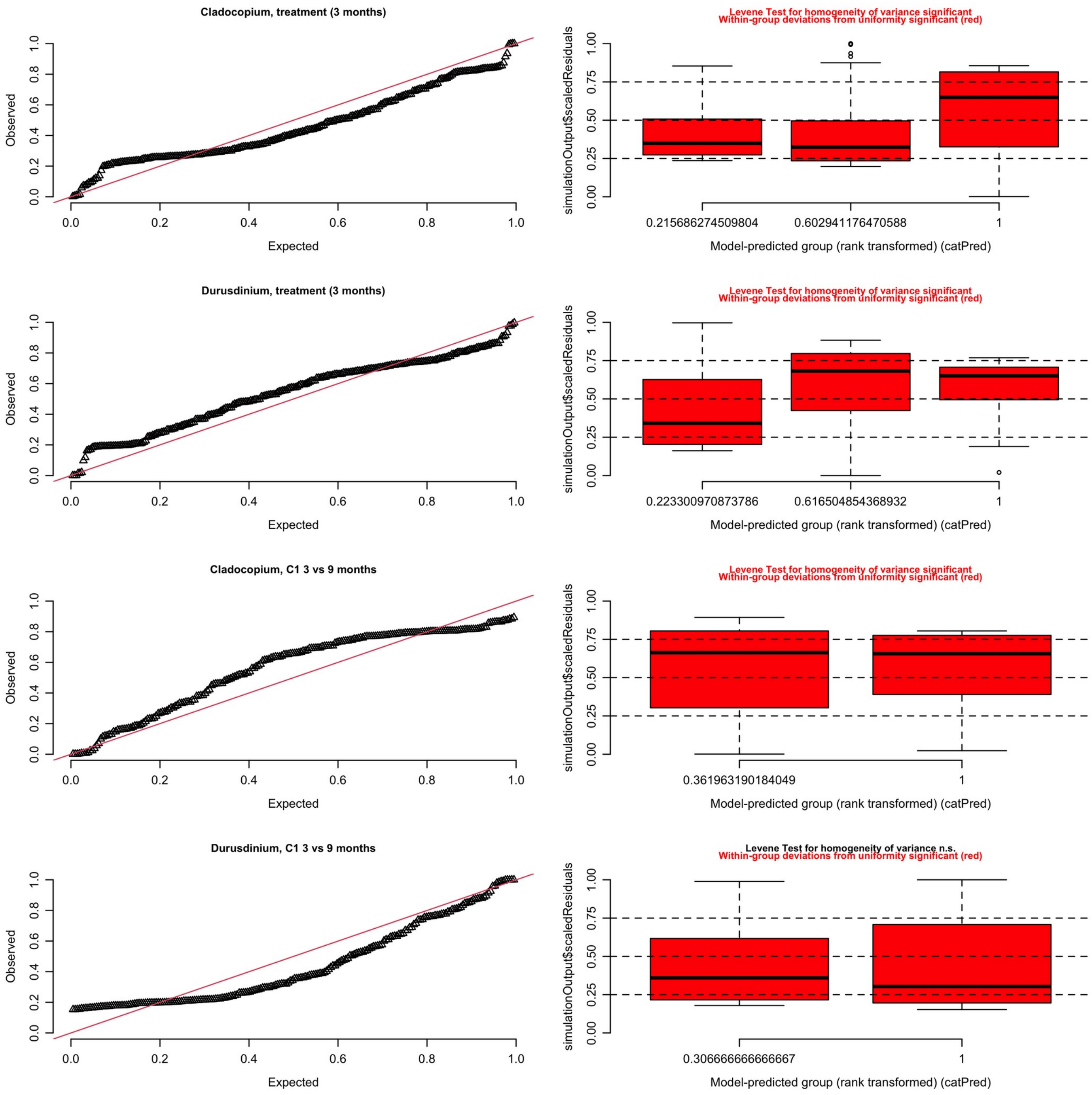

**Table S5.** Detection of fingerprint taxa in samples. SCF049 and SCF086 are the culture codes of the C1 and D1 strains.

**S5A.** Full ranked ITS2 sequence composition of the C1 (SCF049) and D1 (SCF086) culture references, before and after fingerprint threshold filtering. Sequences are ranked by median relative abundance across available culture replicates (one replicate for SCF049; two for SCF086). Cumulative coverage indicates the running total of relative abundance up to and including each sequence; sequences with cumulative coverage ≤ 98% were retained in the culture fingerprint used for downstream field-sample comparisons (see Methods).

***C1 culture (SCF049)***

| Rank | ITS2 sequence | Relative abundance (%) | Cumulative coverage (%) | Replicates present in | In fingerprint (≤98% coverage) |
|---|---|---|---|---|---|
| 1 | C1 | 72.97 | 72.97 | 1 | Yes |
| 2 | C1c | 8.28 | 81.25 | 1 | Yes |
| 3 | C1b | 4.57 | 85.83 | 1 | Yes |
| 4 | C72k | 2.77 | 88.59 | 1 | Yes |
| 5 | C3 | 1.64 | 90.23 | 1 | Yes |
| 6 | C1w | 1.57 | 91.80 | 1 | Yes |
| 7 | C42.2 | 1.36 | 93.16 | 1 | Yes |
| 8 | C3ju | 1.32 | 94.48 | 1 | Yes |
| 9 | C1al | 1.16 | 95.64 | 1 | Yes |
| 10 | C1jx | 0.85 | 96.49 | 1 | Yes |
| 11 | C1bh | 0.71 | 97.20 | 1 | Yes |
| 12 | C3sa | 0.62 | 97.82 | 1 | Yes |
| 13 | C3ew | 0.59 | 98.40 | 1 | No |
| 14 | C42au | 0.57 | 98.98 | 1 | No |
| 15 | C1br | 0.55 | 99.53 | 1 | No |
| 16 | C1t | 0.47 | 100.00 | 1 | No |

***D1 culture (SCF086)***

| Rank | ITS2 sequence | Relative abundance (%) | Cumulative coverage (%) | Replicates present in | In fingerprint (≤98% coverage) |
|---|---|---|---|---|---|
| 1 | D1 | 65.55 | 65.55 | 2 | Yes |
| 2 | D4c | 10.70 | 76.25 | 2 | Yes |
| 3 | D4 | 9.28 | 85.53 | 2 | Yes |
| 4 | D4f | 3.03 | 88.56 | 2 | Yes |
| 5 | D6 | 2.34 | 90.89 | 2 | Yes |
| 6 | D1l | 2.33 | 93.22 | 2 | Yes |
| 7 | D4au | 1.98 | 95.20 | 2 | Yes |
| 8 | D1c | 1.96 | 97.17 | 2 | Yes |
| 9 | D1h | 1.50 | 98.66 | 2 | No |
| 10 | D1hf | 0.82 | 99.48 | 2 | No |
| 11 | 16769_D | 0.52 | 100.00 | 2 | No |

**S5B.** Fingerprint contribution and fingerprint similarity to cultured fingerprint profiles across treatments and timepoints. Fingerprint contribution represents the summed relative abundance of all ITS2 sequences belonging to each culture-defined fingerprint. Mean and median fingerprint similarity represent similarity to the corresponding fingerprint profile, calculated within fingerprint sequence space after renormalisation. High-similarity samples were defined as fingerprint similarity ≥ 0.95.

| Timepoint | Treatment | Culture fingerprint | n samples | Mean fingerprint (%) | Median fingerprint (%) | Mean fingerprint similarity | Median fingerprint similarity | % high similarity |
|---|---|---|---|---|---|---|---|---|
| 3 months | Control | C1 (SCF049) | 71 | 2.54 | 0.00 | 0.01 | 0.00 | 0.00 |
| 3 months | Control | D1 | 71 | 84.02 | 94.75 | 0.87 | 0.95 | 54.93 |

| Timepoint | Treatment | Culture fingerprint | n samples | Mean fingerprint (%) | Median fingerprint (%) | Mean fingerprint similarity | Median fingerprint similarity | % high similarity |
|---|---|---|---|---|---|---|---|---|
| | | (SCF086) | | | | | | |
| 3 months | C1 | C1 (SCF049) | 91 | 69.08 | 95.36 | 0.71 | 0.97 | 63.74 |
| 3 months | C1 | D1 (SCF086) | 91 | 8.09 | 0.73 | 0.21 | 0.01 | 2.20 |
| 3 months | D1 | C1 (SCF049) | 87 | 1.21 | 0.09 | 0.01 | 0.00 | 0.00 |
| 3 months | D1 | D1 (SCF086) | 87 | 94.62 | 96.41 | 0.96 | 0.96 | 82.76 |
| 9 months | C1 | C1 (SCF049) | 117 | 64.91 | 87.26 | 0.68 | 0.91 | 44.44 |
| 9 months | C1 | D1 (SCF086) | 117 | 12.01 | 0.43 | 0.18 | 0.01 | 3.42 |

**Table S6.** Detection frequency and relative abundance of minor non-fingerprint Symbiodiniaceae taxa in recruits across treatments and timepoints. Frequency represents the percentage of samples within each treatment and timepoint in which each taxon was detected. Median relative abundance was calculated only among samples where that taxon was detected.

| Timepoint | Treatment | Taxa | N samples with taxa | Total samples | % samples with taxa | Median relative abundance (%) |
|---|---|---|---|---|---|---|
| 3 months | Control | *Symbiodinium* | 48 | 71 | 67.61 | 0.37 |
| 3 months | Control | *Other Cladocopium* | 64 | 71 | 90.14 | 0.41 |
| 3 months | Control | *Other Durusdinium* | 68 | 71 | 95.77 | 3.38 |

| Timepoint | Treatment | Taxa | N samples with taxa | Total samples | % samples with taxa | Median relative abundance (%) |
|---|---|---|---|---|---|---|
| 3 months | Control | *Fugacium* | 26 | 71 | 36.62 | 0.27 |
| 3 months | C1 | *Symbiodinium* | 46 | 91 | 50.55 | 0.09 |
| 3 months | C1 | *Breviolum* | 1 | 91 | 1.10 | 0.22 |
| 3 months | C1 | *Other Cladocopium* | 91 | 91 | 100.00 | 2.90 |
| 3 months | C1 | *Other Durusdinium* | 81 | 91 | 89.01 | 0.24 |
| 3 months | C1 | *Fugacium* | 8 | 91 | 8.79 | 0.08 |
| 3 months | D1 | *Symbiodinium* | 63 | 87 | 72.41 | 0.07 |
| 3 months | D1 | *Other Cladocopium* | 75 | 87 | 86.21 | 0.23 |
| 3 months | D1 | *Other Durusdinium* | 87 | 87 | 100.00 | 2.74 |
| 3 months | D1 | *Fugacium* | 15 | 87 | 17.24 | 0.03 |
| 9 months | C1 | *Symbiodinium* | 95 | 117 | 81.20 | 0.23 |
| 9 months | C1 | *Other Cladocopium* | 117 | 117 | 100.00 | 3.34 |
| 9 months | C1 | *Other Durusdinium* | 104 | 117 | 88.89 | 0.54 |
| 9 months | C1 | *Fugacium* | 35 | 117 | 29.91 | 0.09 |
| 9 months | C1 | *Gerakladium* | 12 | 117 | 10.26 | 0.04 |
| 9 months | C1 | *Halluxium* | 1 | 117 | 0.85 | 0.01 |
